# Zooplankton impacts on the persistence of the anthropogenic pollution marker *intI*1 in lake water

**DOI:** 10.1101/2024.01.15.575667

**Authors:** Giulia Borgomaneiro, Andrea Di Cesare, Cristiana Callieri, Gianluca Corno, Diego Fontaneto, Roberta Piscia, Ester M. Eckert

**Author notes:** Corresponding author: EM Eckert:.

## Abstract

Wastewater treatment plants (WWTP) effluents can release microbiological pollutants, including the *int*I1 gene (integrases of class 1 integrons), which has been proposed as a target for monitoring anthropogenic pollution in surface waters. This gene has also a strong correlation with antibiotic resistance, making of it an important proxy to evaluate the level of genetic contamination in aquatic environments. he ecological factors that influence the abundance and dynamics of *intI*1 within natural water bodies are largely unknown. To better understand the fate of class 1 integrons in aquatic systems, we resorted to classical limnological monitoring of *intI*1 over multiple years. We also conducted experiments to elucidate the impact of *Daphnia* grazing on its abundance. The monitoring of different size fractions of the Lake Maggiore microbial community has shown a particle-bound life-style for *intI*1-harbouring bacteria. Most of the bacteria hosting *intI*1, originating from both a wastewater effluent that discharges intro Lake Maggiore and lake water itself, grow on particulate substrates in open waters, making them particularly vulnerable to grazing by large filter feeders such as *Daphnia*. *Daphnia* grazing is independent from the origin (lake water or wastewater) of the bacterial genera; it selectively removes bacteria that are present in aggregates or even filamentous forms from both origins. To understand if *intI*1 is related to viable bacteria or just DNA residues, it is important to study the persistence of class 1 integrons with their gene cassettes, which often contain antibiotic resistance genes in freshwater ecosystems.

**Significance Statement:** While faecal pollution of freshwaters is commonly monitored, genetic pollution through wastewater treatment plant outflows, such as antibiotic resistance genes, is difficult to monitor due to the diverse nature of genes present. The *intI*1 gene is proposed as a proxy for anthropogenic pollution; however, there is a major lack of understanding regarding the persistence of this gene in freshwaters. In this study, we demonstrate that *intI*1 in freshwaters is associated with both the natural microbial community and allochthonous microbes arriving from wastewater. Furthermore, we show that *intI*1 harbouring bacteria preferentially reside in the aggregated microbial fraction and are easily removed by zooplankton grazing. This study is the first limnological investigation of this gene and highlights a significant gap in our knowledge regarding the ecology of class 1 integrons.

Genetic pollution of surface waters is however a global problem and of very broad interest on the one hand, on the other hand, the question of the establishment of an allochthonous gene into a natural microbial community is also an interesting fundamental question in ecology, thus this study has both more applied and more fundamental aspects. Therefore, we consider it perfect for the readership of L&O.

## Introduction

Monitoring is required to evaluate and manage the impact of anthropogenic pollution in aquatic environments, and thanks to low costs and easy reproducibility, DNA-based methods are considered the future for the field (Zolkefli et al. 2020). One of the most important polluters of surface waters in industrialised countries is the effluent of wastewater treatment plants (WWTP), which can release fecal bacteria (Marano et al. 2020), and is a source of other microbiological pollutants, including genetic pollution by *i.e.*, antibiotic resistance genes (ARGs) (Munk et al. 2022). Several genes have been suggested as potential monitoring targets for this kind of pollution. Among these, the frequency of the *intI*1 gene exhibits a strong correlation not only with anthropogenic pollution in general but also with the antibiotic resistance specifically, making it one of the most promising targets for DNA-based environmental monitoring (Gillings et al. 2015; Zheng et al. 2020; Abramova et al. 2022). Integrons are genetic elements that capture and express genes, and the *intI* genes encode for integron integrases, proteins that catalyse the recombination of exogenous gene cassettes with a specific recombination site (Gillings 2014). Therefore, integrons are important in the acquisition and dissemination of ARGs. Thanks to its strong correlation with anthropogenic pollution, quantifying *intI*1 in microbial community DNA extracts from water samples is a potent and uprising method to evaluate the level of genetic contamination in aquatic environments (Zheng et al. 2020).

The *intI*1 gene is generally found in freshwater ecosystems even when they are exposed to low anthropogenic impact (Cacace et al. 2019; Abramova et al. 2022). However, due to its correlation with anthropogenic pollution some authors have gone as far as to calling the class 1 integrons an “invasive species” (Gillings 2017). The term “invasive species” has many contradictory definitions, but a shared criterion is the spread of the invader within the invaded ecosystem (Pereyra 2016). Whether allochthonous *intI*1 is indeed able to spread within lake ecosystems is unknown, since questions regarding stability and dynamics of *intI*1 within natural waterbodies have been addressed only by a limited number of studies (Lee et al. 2021). Physical and chemical parameters that might be considered intrinsic properties of water bodies such as temperature or pH might influence the abundance of *intI*1 within the system. Yet, the few data available is still contradictory (Di Cesare et al. 2020; Chaturvedi et al. 2021). Generally, it seems that the quantity of *intI*1 can be linked to the proportion of WWTP effluents entering freshwaters, suggesting a continue external introduction, rather than a spread by growth of bacteria hosting class 1 integrons (Haenelt et al. 2023; Corno et al. 2023), contrary to its definition as an “invasive species”. In terms of water quality, *intI*1 abundances seem to readily increase in experimental communities with the addition of even very low quantities of wastewaters treated at the highest standards (Subirats et al. 2019) and the stronger the contamination the more abundant the class 1 integrons seem to be in natural microbial communities (Di Cesare et al. 2022b). In fact, so far only two studies have observed an increase of the abundance of the gene without apparent chemical water pollution or direct seeding, suggesting growth of *intI*1 containing bacteria within the natural water body (Lee et al. 2021; Haenelt et al. 2023).

Which ecological factors influence the abundance of *intI*1 is largely unknown. The integrons themselves are not mobile genetic elements, and *intI*1 is usually encoded in the chromosome (Gillings 2014). Thus, its fate in freshwater systems, for long-term persistence, is largely determined by the ecological success of the bacterial carrier. Abiotic factors such as low nutrient availability and biotic factors such as competition with the autochthonous bacterial community and predation are important variables that reduce the ecological success of many allochthonous bacteria entering the lake from WWTP outflows (Korajkic et al. 2019). This is mainly attributed to the fact that such genotypes are adapted to fast growth in high nutrient conditions, which makes of them on the one hand poor competitors in oligotrophic systems and on the other hand a preferential prey for heterotrophic nanoflagellates. Given that *intI*1 can reside in various bacterial hosts across different taxa, it remains to be tested which factors could influence the overall abundance of this gene (Lee et al. 2021).

To illustrate the potential complexity of the interactions and how they might influence *intI*1 dynamics one could make the following considerations: in a study in Lake Maggiore, we found that the gene was enriched in the sediment compared to the water column, suggesting that there was a general trend of the gene sinking down with particulate organic matter from surface waters and accumulating in depth, with close to no resuspension from the sediment (Di Cesare et al. 2020). This would imply that the bacteria hosting *intI*1 are less commonly free-living but might rather resort to a particle-associated live-style (Di Cesare et al. 2020). This aggregation or attachment might protect *intI*1 harbouring bacteria from small flagellated grazers (Corno and Jürgens 2008) but might make them more vulnerable to filter feeders, such as *Daphnia* or certain rotifers since they preferentially feed on larger particles (Langenheder and Jürgens 2001).

In order to better understand the fate of class 1 integrons in aquatic systems we resorted to classical limnological monitoring of the *intI*1 gene over multiple years in three different size fractions, namely: (1) free-living bacteria, (2) particle-attached bacteria, and (3) zooplankton-attached bacteria. Then, we conducted experiments with natural bacterial communities with model organisms from multiple trophic levels, such as the crustacean *Daphnia obtusa*, the flagellate *Poteriochromonas* sp, and the rotifer *Rotaria macrura*. Since *Daphnia* seemed to impact the abundances of *intI*1, experiments to elucidate the role of *Daphnia* grazing on its abundance were further conducted. Thereby we tried to disentangle the effect of *Daphnia* on microbes deriving from a WWTP effluent and its allochthonous *intI*1 containing bacteria and on *intI*1 hosting bacteria that are naturally found within the lake ecosystem.

## Materials and Methods

### Sampling and filtration

Lake Maggiore is a large subalpine lake shared between Italy and Switzerland. The water samples were taken monthly from January 2019 to December 2021 at two sampling sites: Ghiffa (WGS84 coordinates: 45°5611N, 8°3811E) and Pallanza (45°5511N, 8°3211E), the former considered a pelagic site and the second a semi-littoral site. The water samples from the lake were collected in a single operation by a sampler that collects a 5-L integrated sample from the thermocline depth up to the surface (Bertoni et al. 2010). Water samples were stored at 4 °C in the dark until processing within 24 h. The natural bacterial community was pre-filtered over 10 μm and filtered on 0.2 μm polycarbonate filters (47 mm diameter) to obtain the free-living fraction of the bacterial community (1). The particle-associated fraction (2) was recovered by filtering the water samples on GFC filters (porosity around 1.2 μm, 47 mm diameter). Zooplankton organisms were collected monthly from January 2019 to December 2021 in Lake Maggiore only in the deepest zone of the lake (Ghiffa station). Samples were collected using a nylon net of 50 µm mesh, 0–50 m depth vertical hauls, in order to reach 1 g d.w. for each sample. Zooplankton and their attached bacteria were collected from the net and these samples were stored in a TRIS/EDTA/NaCl (each 0.1 M) buffer and frozen at – 20°C until analyses. They represent the zooplankton-attached fraction (3) of the bacterial community.

In order to relate gene abundances to the physical and chemical characteristics of the lake a dataset including NO_3,_ TN, Si, NH_4_, Ca, Mg, Na, Cl and SO_4_ with monthly samples from Lake Maggiore was used . Organic carbon concentration was measured using a total organic carbon (TOC) analyzer while *in-situ* measurements of temperature, conductivity, pH, and dissolved oxygen were carried out using a multiparametric probe (CTD316, Idronaut, Brugherio, Italy), (Callieri et al. 2022).

Differences in both the occurrence and abundance of *intI*1 across the three filtration fractions, sampling year, and sampling locations were tested using linear models in R version 4.2.2 (R Core Team 2022) in R-Studio. Model fit was evaluated using the *performance* package (Lüdecke et al. 2021). Interactions between the tested variables were removed from the models when they were not significant. For significant categorical predictors with more than two levels, post-hoc tests were performed by estimated marginal means (EMMs) using the *emmeans* package (Lenth R. 2023).

In order to analyse the influence of physical and chemical parameters on the abundance of the gene further models were made for the years and fractions with a high number of quantifiable samples, namely 2020 and 2021 and the fractions of free-living bacteria (0.2 μm filters) and particle-attached bacteria (GFP filters). To reduces the variables to be used in the model they were first correlated via Pearson’s moment correlation and variables with a r ≥ 0.8 were considered correlated (Tabel S1) and consequently the following variables were chosen to be integrated in the model: Temperature (negatively correlated to NO_3_, TN and Si), O_2_ (negatively correlated to Si and positively to pH and NH_4_), Conductivity (positively correlated to Alkalinity, Ca, Mg, Na, Cl, SO_4_), TP and TOC. A linear model was constructed using log transformed data for gene abundances. Model fit was evaluated with the *performance* package. Plots were made for variables that significantly influenced the gene abundances using *ggplot2* (Wickham H. 2016) and *cowplot* (Claus O. Wilke 2020).

### Food web experiment and analysis

The experiment was performed to test the effect of predators of bacteria in the food web of Lake Maggiore, by adding different predators to a natural bacterial community containing *intI*1 (figure 1). The predators were chosen from the filter feeder crustacean *Daphnia obtusa* (with densities of 1 adult animal in 100ml), the filter feeder bdelloid rotifer *Rotaria macrura* (1 animals in 10 ml), and the mixotrophic nanoflagellate *Poteriochromonas* sp. (10^3^ cells ml^-1^). These organisms were chosen to study the impact grazing on bacteria, as previously investigated by (Ricci 1984; De Bernardi et al. 1985; Corno and Jürgens 2006). All possible combinations of one, two, and three predators were added in flasks with 100 ml of Lake Maggiore water pre-filtered over 5 μm with a microbial abundance of 10 cells ml . Surface water samples from Lake Maggiore were taken manually from the shore. Each combination was conducted in triplicate. The treatments were incubated for 72 h then the whole community was filtered on 0.2 μm polycarbonate filters (25 mm diameter). Samples were stored in TRIS-EDTA buffer and frozen at – 20°C until further molecular analyses (quantification of the *intI*1 and 16S rRNA genes by quantitative Real Time PCR, qPCR). Flagellate and bacterial abundances at the beginning of the experiment were determined by the flow c Accuri C6 (Becton Dickinson, Oxford, UK), equipped with a 20 mW 488 nm Solid State Blue Laser and a 14.7 mW 640 nm Diode Red Laser.

**Figure 1:**
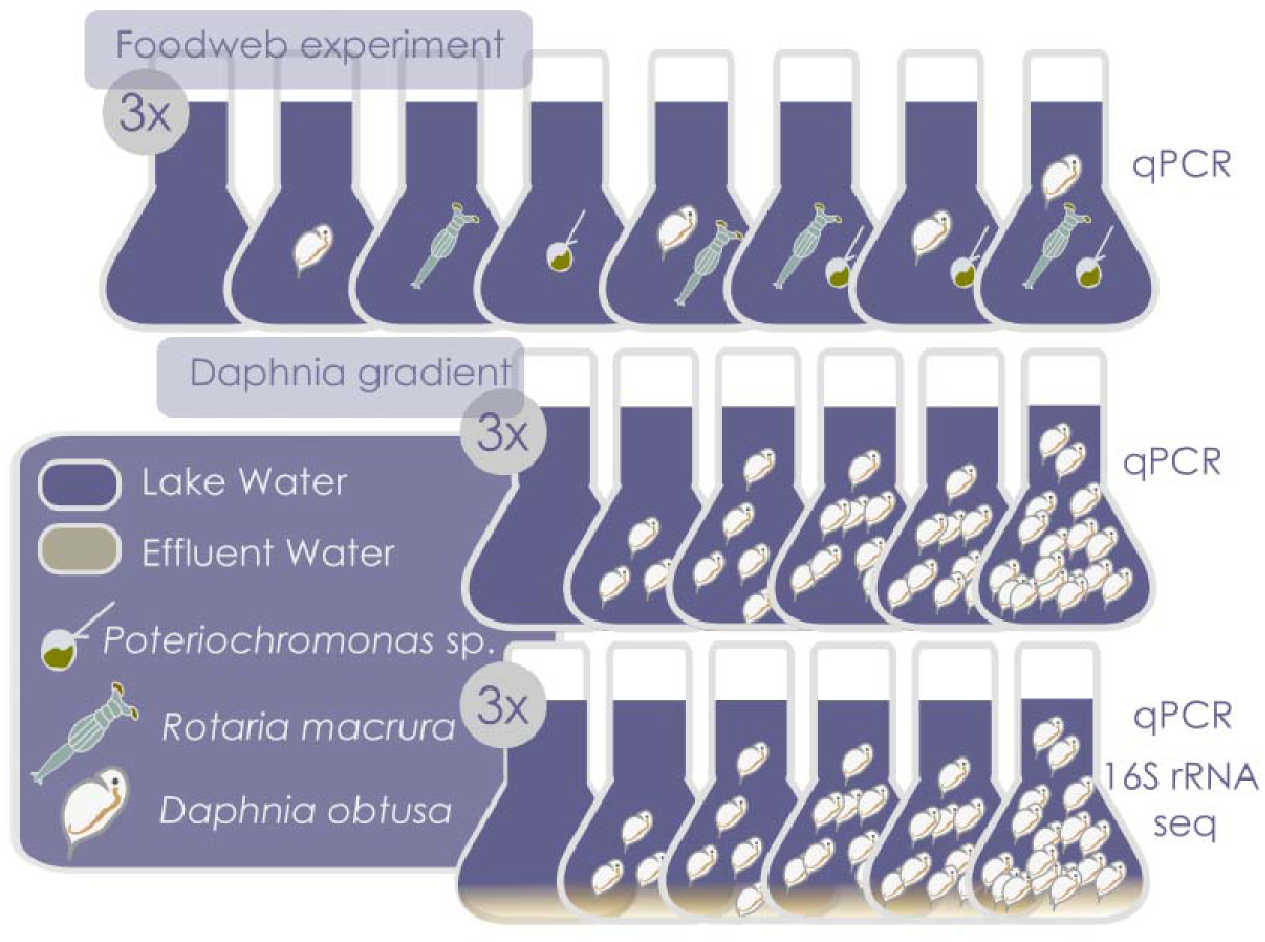
Schematic description of the experimental set up of the three experiments (foodweb: *Daphnia* in lake water, *Daphnia* in lake water with WWTP effluent addition) conducted in the study and the analysis conducted on the extracted DNA.

A linear model was conducted to test the effect on (log transformed abundance of) *int*I1 of the presence of the different predators in the food web and their interaction. Model quality was evaluated using the *performance* package. Plots were made using the packages *ggplot2* and *cowplot* with a logarithmic scale for the gene abundance. Interactions were omitted from the final model if not significant.

### *Daphnia* gradient experiment and analysis

Two experiments were performed, using different water matrices: the first one with lake water and the second one with lake water and WWTP effluent water. This was done in order to see whether *Daphnia* had a distinct effect on bacteria deriving from WWTP effluent or natural *int*I1 containing bacteria (figure 1).

i. Lake water. A sample of Lake Maggiore surface water was taken manually from the shore. Samples of *D. obtusa* derived from a long-term culture, kept in a small garden pond at the CNR-IRSA in Verbania. Animals were collected from the garden and cultured under laboratory conditions up to the second generation to avoid maternal effect (Ebert 1993; LaMontagne and McCauley 2001; Lampert 2001); these laboratory-adapted animals were used in the experiments*. Daphnia obtusa* was added to the flask, with 100 ml of lake water and a quantity of animals in a gradient with the following numbers of animals per vessel: 0, 3, 6, 9, 12, and 15. Each treatment was conducted in triplicate. The treatments were incubated for 72 h at 22°C in the dark. The bacterial community at the end of the experiment was pre-filtered over 25 μm to remove the animals, filtered on 0.2 μm polycarbonate filters (25 mm diameter) and kept frozen at − 20°C for quantification of the *intI*1 and 16S rRNA genes by qPCR.
ii. Lake water with 10% WWTP effluent water. Experiment II was essentially performed as described for experiment I, except that we used a mix of lake water with 10% WWTP effluent water. The effluent water samples were collected at the large municipal WWTP of Verbania (45°56’, 8°33’; 51,000 Population Equivalent, PE in Eastern Piedmont, Northern Italy). This WWTP receives domestic sewage and an amount of pretreated hospital sewages. These samples were used for both qPCR (to quantify the *intI*1 and 16S rRNA genes) and microbial community analysis (by 16S rRNA gene amplicon sequencing).

For both experiments the relationship between the abundance of *int*I1 and number of animals was tested in a linear model. Model quality was visually inspected by plotting the residuals of the model. Plots were made using the packages *ggplot2* and *cowplot* with a logarithmic scale for the gene abundance.

### DNA extraction and qPCR analysis

The filters were processed to extract DNA by using a commercial kit (DNeasy UltraClean Microbial Kit, QIAGEN) according to the manufacturer’s instructions with the modifications mentioned in (Di Cesare et al. 2015). The zooplankton organisms were preserved in a TRIS/EDTA/NaCl (each 0.1 M) buffer, an aliquot (0,5 ml) of these bulk samples were placed in the PowerBead tubes of the PowerSoil DNA extraction kit (Qiagen) and supplemented with 1% SDS and 250 µg ml^-1^ of Proteinase K at -20 °C until DNA extraction. The samples were homogenized using the Precellys instrument in two cycles, each at 6000 rpm for 1 minute. Subsequently, they were incubated at 52 °C for 2 hours. Then, each incubated sample was processed for the DNA extraction following the manufacture’s instruction of the PowerSoil DNA extraction kit (Qiagen).

The DNA extracts were tested by qPCR for the abundance of the class 1 integron integrase (*intI*1) and the 16S rRNA genes. The *intI*1 gene abundance was normalized to the abundance of the 16S rRNA gene. All qPCRs were performed using the RT-thermocycler CFX Connect (Bio-Rad, Italy). The primer pairs used to quantify the 16S rRNA gene were Bact1369 F 511CGGTGAATACGTTCYCGG311 and Prok1492 R 511GGHTACCTTGTTACGACTT3’ and to quantify the and the *int*I1 gene were intI1LC5 511GATCGGTCGAATGCGTGT311 and intI1 LC1 511GCCTTGATGTTACCCGAGAG3’ (Barraud et al. 2010; Di Cesare et al. 2015). The protocols and programs used to quantify the 16S rRNA and *intI*1 genes were previously described in (Di Cesare et al. 2015) and (Di Cesare et al. 2016) respectively. The standard curve and inhibition tests were performed as previously described (Di Cesare et al. 2013) and no inhibition was determined. The efficiency of reactions, R^2^, and the limits of quantification were determined as previously described (Bustin et al. 2009). The mean value ± standard deviation of the efficiency of reactions and R^2^ were 97.17 ± 6.73 and 0.99 ± 0.01, respectively. The limits of quantification (LOQ) are shown in Table S2. The correct size of all qPCR products was evaluated by electrophoresis (30 min at 80 V, 1% agarose gel).

### 16S rRNA gene sequencing and analysis

DNA samples from the second experiment, where *int*I1 was quantified in the microbial community from both lake and effluent water (9:1) by qPCR, were additionally used for amplicon sequencing of the V3/V4 of the 16S rRNA gene. Using the universal bacterial primers S-D-Bact-0341-b-S-17 (forward primer, 511-CCTACGGGNGGCWGCAG-311) and S-D-Bact-0785-a-A-21 (reverse primer, 511-GACTACHVGGGTATCTAATCC-311) (Herlemann et al. 2011), sequencing was conducted on an Illumina MiSeq platform by Macrogen (Seoul, South Korea). The raw amplicon reads of the bacterial community are available in the NCBI database and can be accessed under project ID PRJNA962832. Data processing was performed in R version 4.2.2. Raw reads were processed using the DADA2 package version 1.26 (Callahan et al. 2016). Additional packages that were used to analyse the data and data handling include phyloseq package version 1.42.0 (McMurdie and Holmes 2013), Biostrings package version 2.66.0 (Pagès et al. 2023) and packages *ggplot2*. The DADA2 pipeline provided in the online tutorial (https://benjjneb.github.io/dada2/tutorial.html) was used with only minor adjustments. Briefly, fastq files were quality checked, filtered (maxEE 2 and 2), and trimmed (left trim 50 bp for R1 and R2 and right trim 25 bp for R2). The obtained sequences were merged into a unique sequence and, through error correction, Amplicon Sequence Variants (ASVs) were determined. They were ten checked for chimeric sequences and used for the taxonomic assignment through the SILVA database Version 138.1 (Quast et al. 2013). ASVs that were not identified at least at the level of phylum or that were identified as either Archaea, chloroplast, or mitochondria were removed from the dataset. The original dataset was composed of 4498 ASVs and 4310 ASVs were retained after processing. Beta-divesity was evaluated using both abundance and occurrence based Bray-Curtis dissimilarity with the *vegan* package and an n*on-metric multidimensional scaling* (nmds) was calculated and plotted with the *phyloseq* package. In order to evaluate the impact on *Daphnia* grazing on the removal of WWTP-derived microbiota compared to the lake microbiota we calculated the distance of each microcosm community to the original lake water and WWTP effluent samples, respectively, and plotted the data per number of *Daphnia*.

Read numbers of genera were correlated to the abundance of *Daphnia obtusa* in all mesocosm samples using Spearman correlation and genera that correlated negatively with *Daphnia* abundance (r>0.8) were used for further analysis. We then categorised bacterial genera in “freshwater derived” or “WWTP derived” or common to both types of water “neutral” using AncomBC analysis (Lin et al. 2022) . Thus we plotted all genera that were significantly negatively correlated to the abundance of *Daphnia obtusa* and checked whether they were more commonly related to WWTP effluent or lake water.

## Results

### **intI**1 dynamics in Lake Maggiore

The *intI*1 gene was detected in 99% of the samples of free-living bacteria (water fraction filtered at 0.2 µm), a significantly higher occurrence compared to the other two fractions where it was detected in 61% (particle-associated bacteria, >1.2 µm) and 32% (zooplankton-associated bacteria, >50 µm) (Table 1A). Despite being detected in fewer samples, normalised copy numbers of the *int*I1 gene per 16S rRNA gene were significantly higher in the samples from the particle-associated bacteria compared to those from the free-living bacteria (Table 1D&E). The mean abundance of *int*I1 was two to three orders of magnitudes higher in particle-attached bacteria compared to the free-living fraction. No differences were found for the gene occurrence or quantity for the two different sampling stations (Table 1B&D). *int*I1 was quantifiable only in 8 samples from the zooplankton-associated fraction, but, when quantifiable, its normalised abundances were notably high (Table 1 & Figure 2A). Despite the lower number of quantifiable samples in the free-living fraction of 2019, the abundances of the *intI*1 gene did not show significant changes over the years (Figure 2B). The gene abundance was only slightly higher in the first months of the year in the free-living fraction, and this was even more pronounced in the first seven months of the year for both the particle-associated and zooplankton-associated fractions (Figure 2B).

**Table 1:**
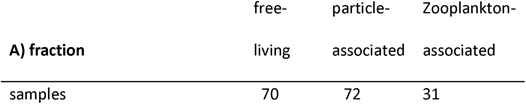
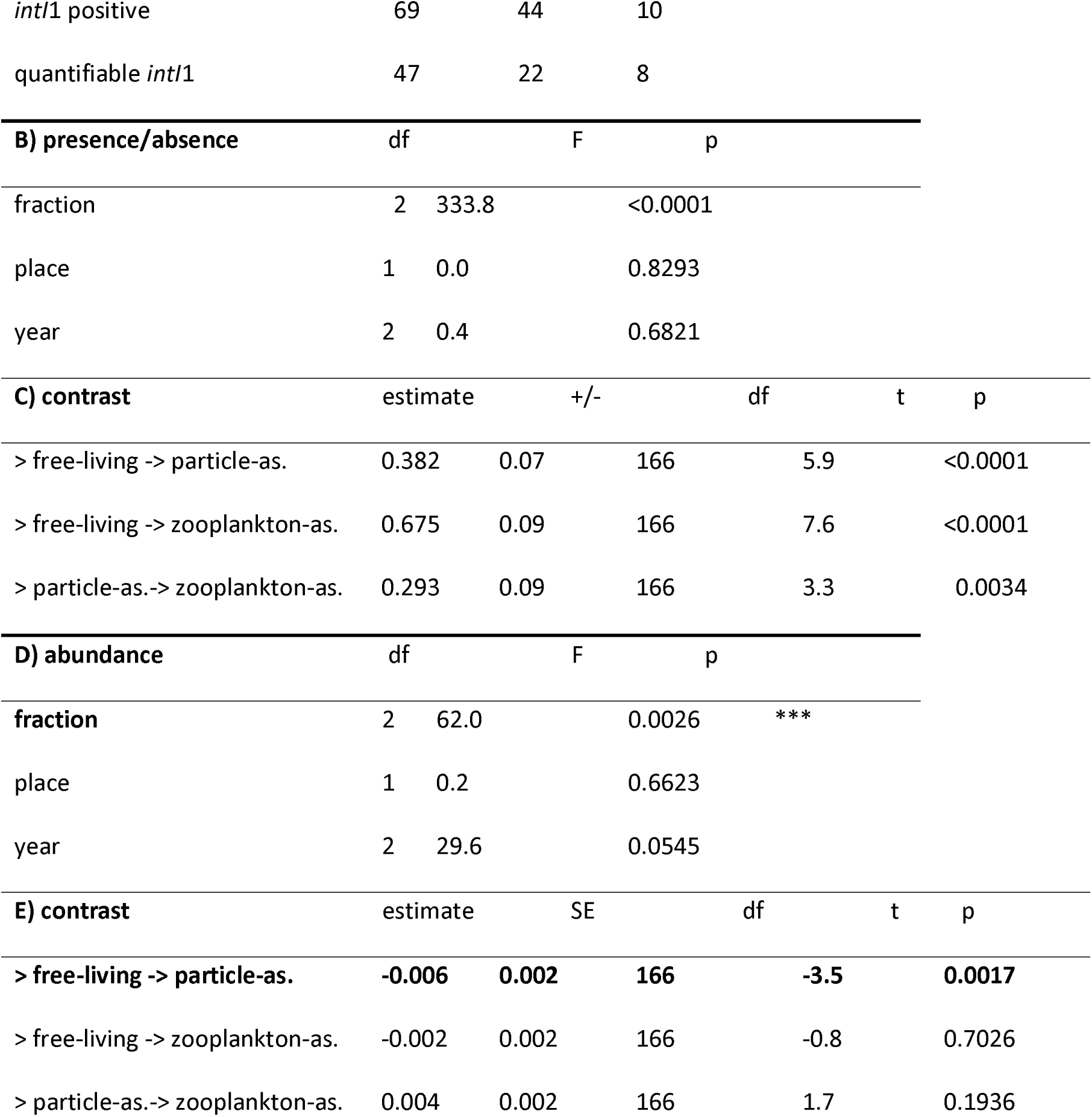
A) Number of samples tested for each fraction, number where the gene *intI*1 was found and number of samples where the gene was quantifiable (quantities depicted in Figure 2). B-D) Output from the linear models testing the effect of fraction and sampling location (place) and year on the occurrence of *int*I1 (presence/absence, B) and the abundance (D) and the post-hoc test between the three tested fractions (C&E).

**Table 2:**
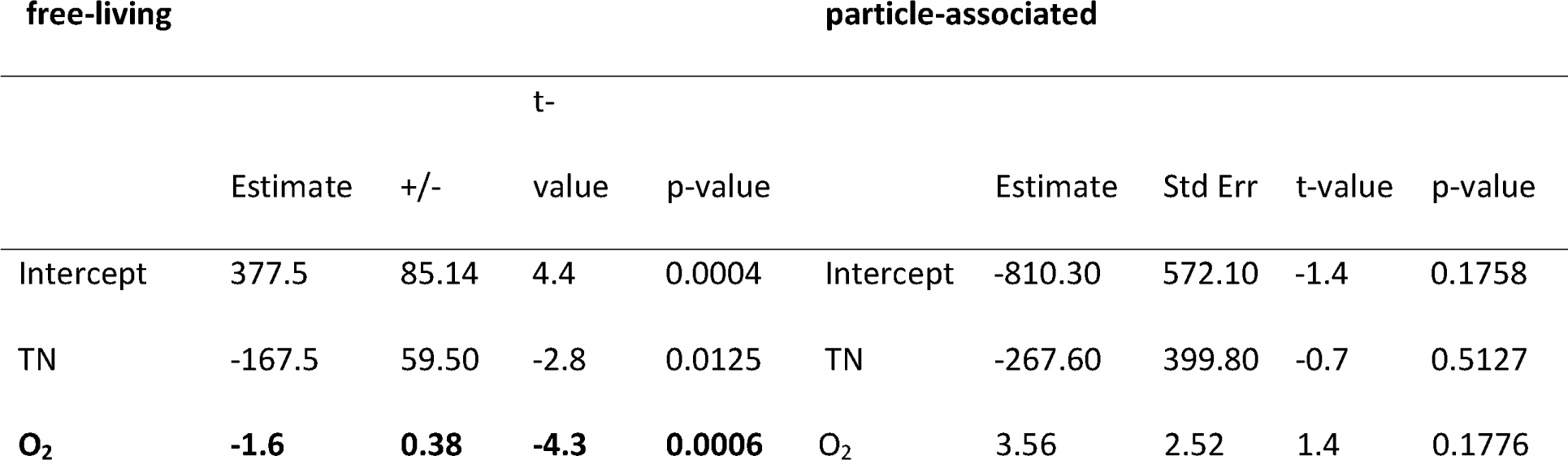
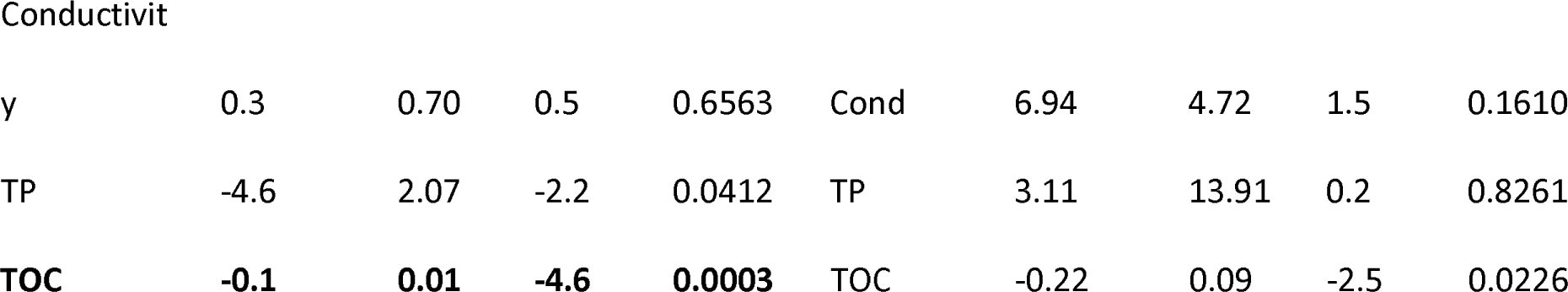
Output of the linear model of the effect of the abundance of *intI*1 in the 0.2 and 1.2 µm fraction in the years 2020 and 2021 with the concentration of total nitrogen (TN), saturation of % Oxygen (O_2_), conductivity, total phosphorus (TP) and total organic carbon (TOC). Significant predictors are marked in bold.

**Figure 2:**
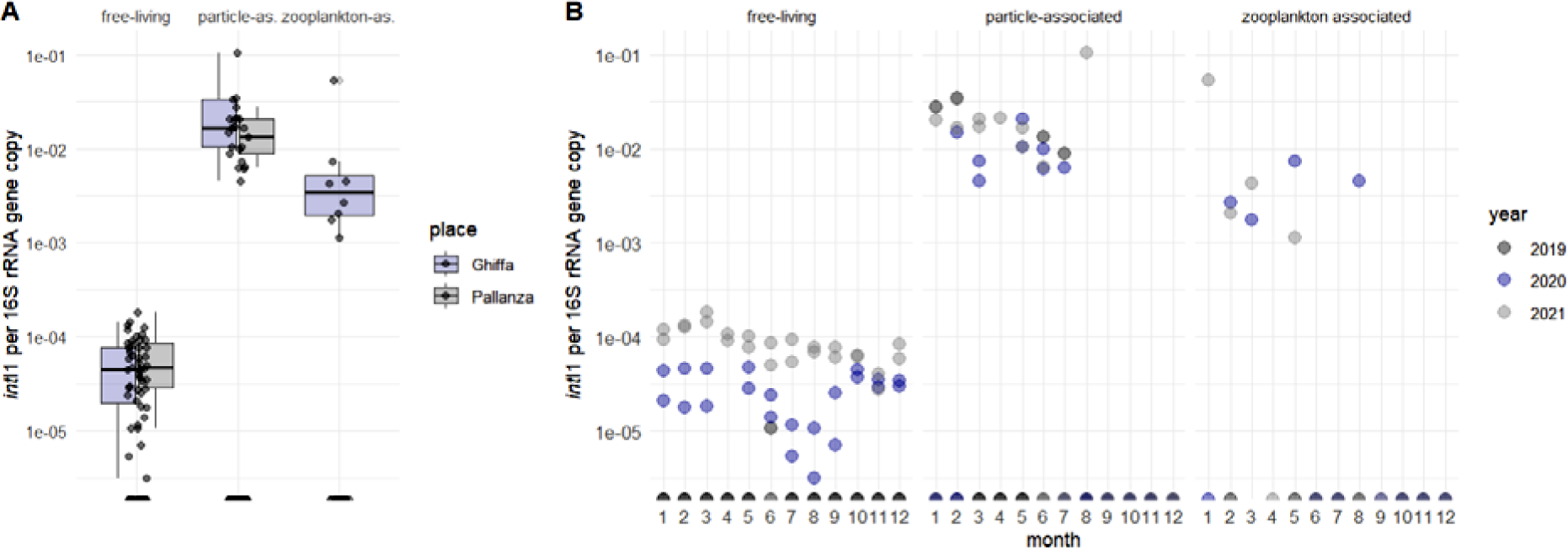
A) Boxplots of normalised abundance of the *int*I1 gene in three different water fractions of samples with quantifiable *int*I1 abundances of free-living and particle-associated at both stations, Ghiffa and Pallanza, and in the zooplankton-associated fraction, Ghiffa station only. B) Dotplots of the same data plotted per fraction and per month of sampling in three different years. Considering the chemical parameters of the lake, the abundances of *intI*1 (years 2020 and 2021) were strongly negatively affected to the saturation of the water with O_2_ and to TOC content (Table 2), both of which were clearly higher in summer and towards the end of the year (Figure S1).

### Food web impact on **intI**1 abundance

By adding filter feeders, the crustacean *Daphnia obtusa* and/or the bdelloid rotifer *Rotaria macrura*, and/or *Poterioochromonas* sp. in all possible combinations, we constructed artificial food webs in microcosm experiments and tested their impact on the abundance of *intI*1. Statistical analysis showed that only the addition of *Daphnia* had a significant negative effect on the abundance of *intI*1 in the system (Table 3 and Figure 3).

**Figure 3.**
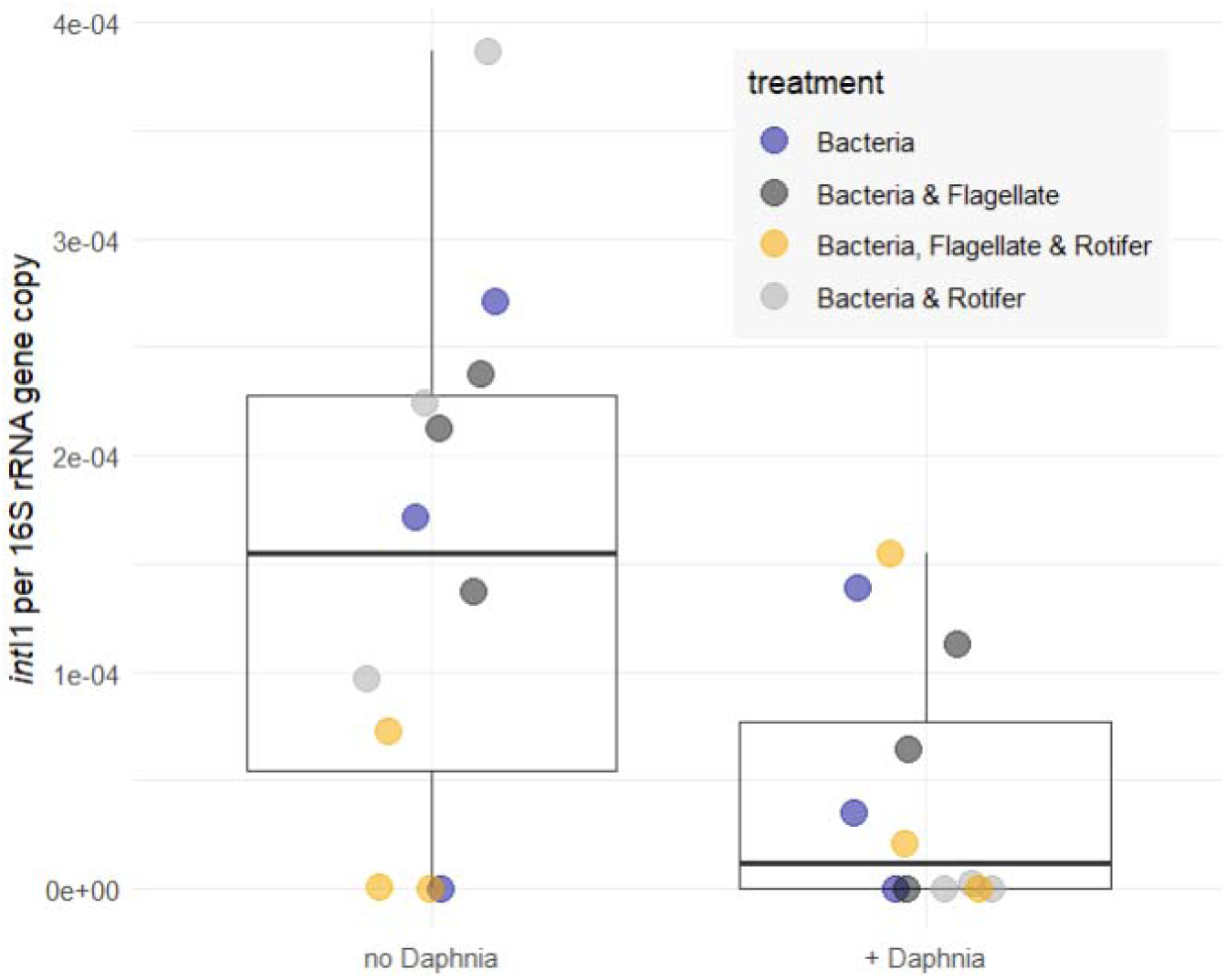
Normalised abundance of the *intI*1 gene in experimental microcosms with and without the addition of *Daphnia obtusa*. Colours indicate the presence of flagellates and rotifers alone and together in the microcosms.

**Table 3.**
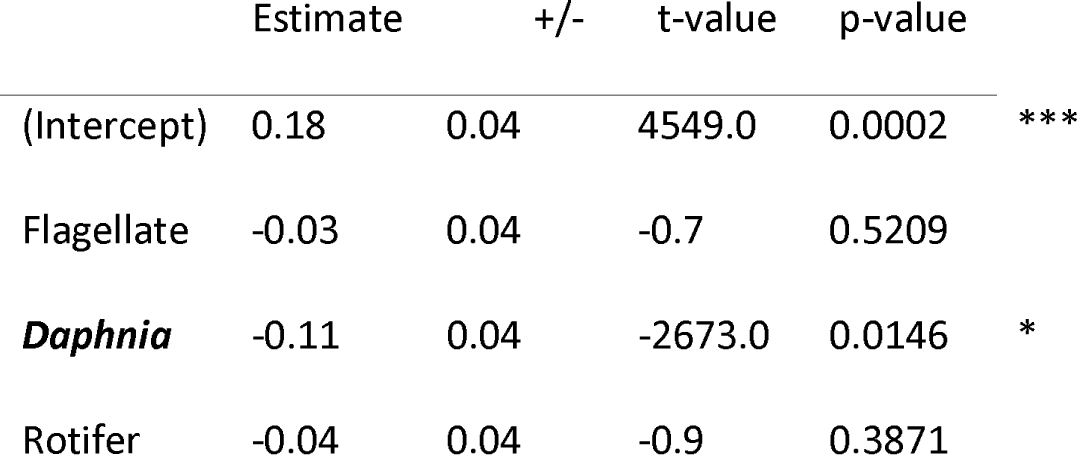
Output from the linear model testing the effect of the various trophic levels on the abundance of *int*I1 in the microcosms. Flagellate=*Poterioochromonas* sp., *Daphnia*=*Daphnia obtusa* , Rotifer=*Rotaria macrura*

### Impact of Daphnia grazing on microbial community

In order to quantify the impact of *Daphnia* on *intI*1 abundance, we conducted two experiments with a gradient of *Daphnia*, one with only lake water and one with the addition of 10% WWTP effluent. In all the samples, *intI*1 was positive and quantifiable. In the microcosms with 100% lake water, the overall abundance of *intI*1 per sample ranged from 8.8 x 10^-3^ to 4.2 x 10^-2^ *intI*1 per copy of 16SrDNA (Figure 4).

**Figure 4.**
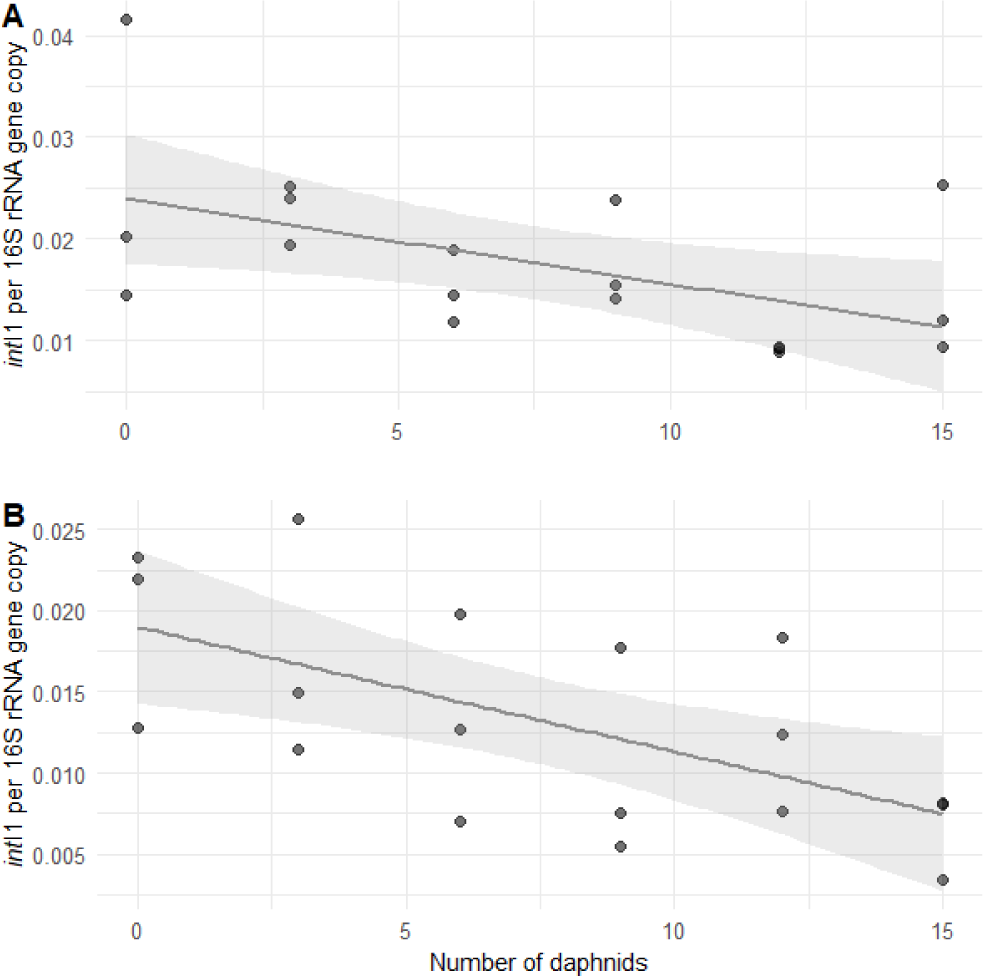
Normalised *int*I1 abundances in microcosms per number of *Daphnia* added to 100ml for: A) only lake water and B) lake water with 10% WWTP effluent. Grey line and shaded area is the trend line and the confidence interval of the linear model.

Instead, in the microcosms with lake water with 10% WWTP effluent, its concentration ranged from 3.4 x 10^-3^ to 2.6 x 10^-2^ gene copies per 16SrRNA gene copy. The abundance of the *intI*1 gene decreased significantly in the water when the number of *Daphnia* increased in both treatments (Figure 4, Table 4).

**Table 4.**
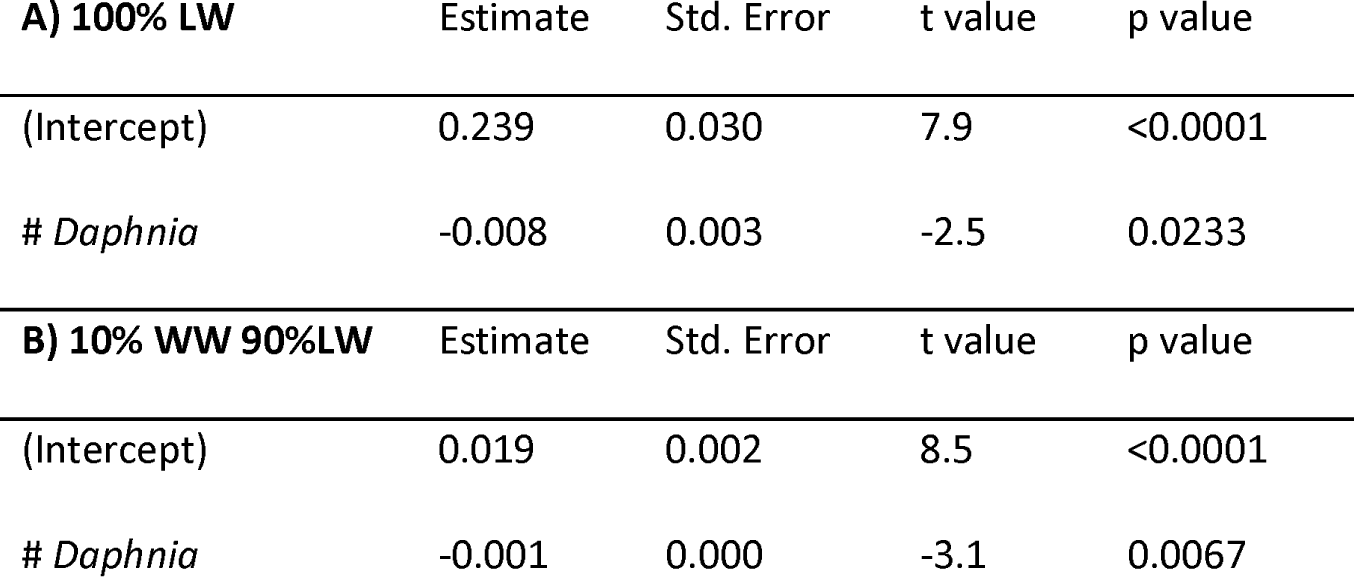
Results from the linear models testing the effect of the number of *Daphnia* on the abundance of the *int*I1 gene in the experiment with 100% lake water (LW) or 10% WWTP effluent water (WW) and 90% LW

The differences in microbial community composition (Beta-diversity) were measured as Bray-Curtis dissimilarity in the WWTP effluent, in the lake water before the experiment and in the water with increasing numbers of *Daphnia* (Figure 5A). Samples from WWTP effluent and in the lake water formed separated clusters. Even the samples with 10% of WWTP effluent without *Daphnia* formed a separated cluster (Figure 5A). In the samples with *Daphnia* (D), the variability of the composition of the microbial community between replicates increased.

**Figure 5:**
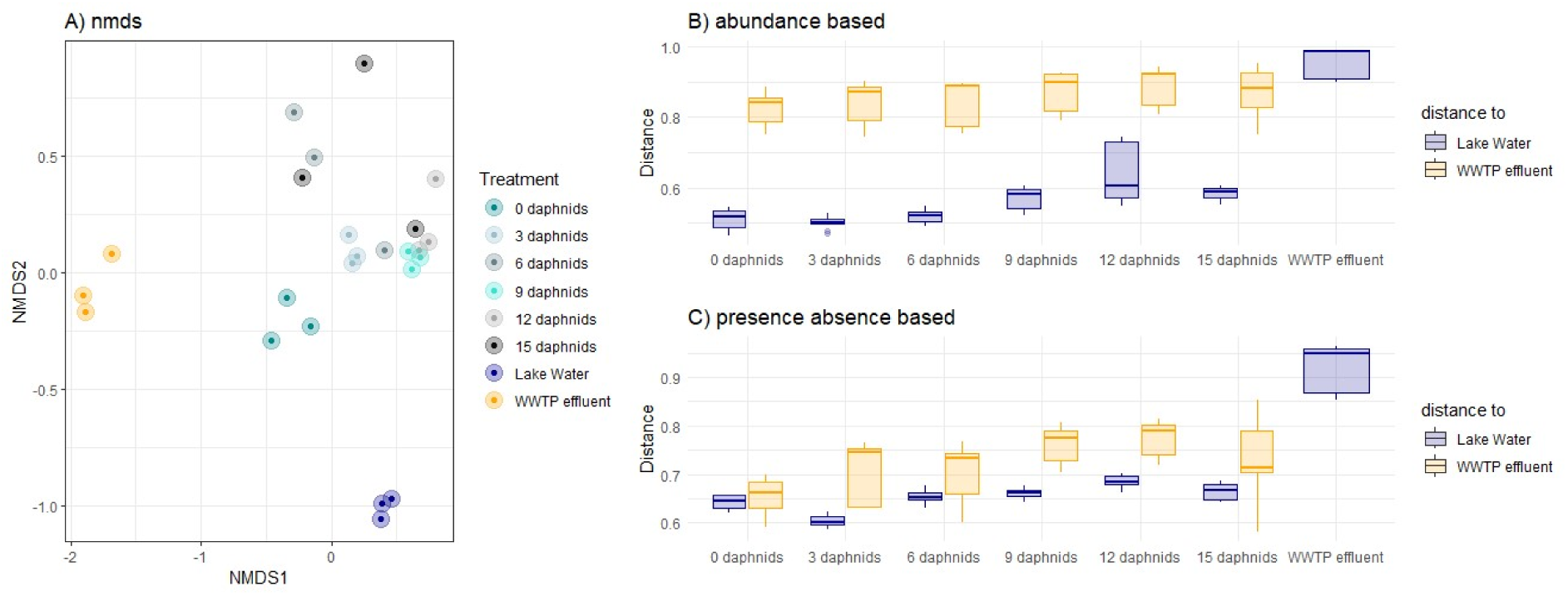
Beta-diversity of the microbial community of each microcosm (denoted by the number of daphnids in 100ml of microcosm) A) nmds plot of abundance-based Bray-Curtis distances, B & C distance to either the original lake water or original WWTP effluent water community calculated with B) abundance and C) occurrence based Bray-Curtis index (β-diversity).

The distance to both Lake Water and WWTP community increased significantly with the increase of *Daphnia*, for both abundance-based and presence absence-based Bray-Curtis distances (Table 5, Figure 5B & C).

**Table 5:**
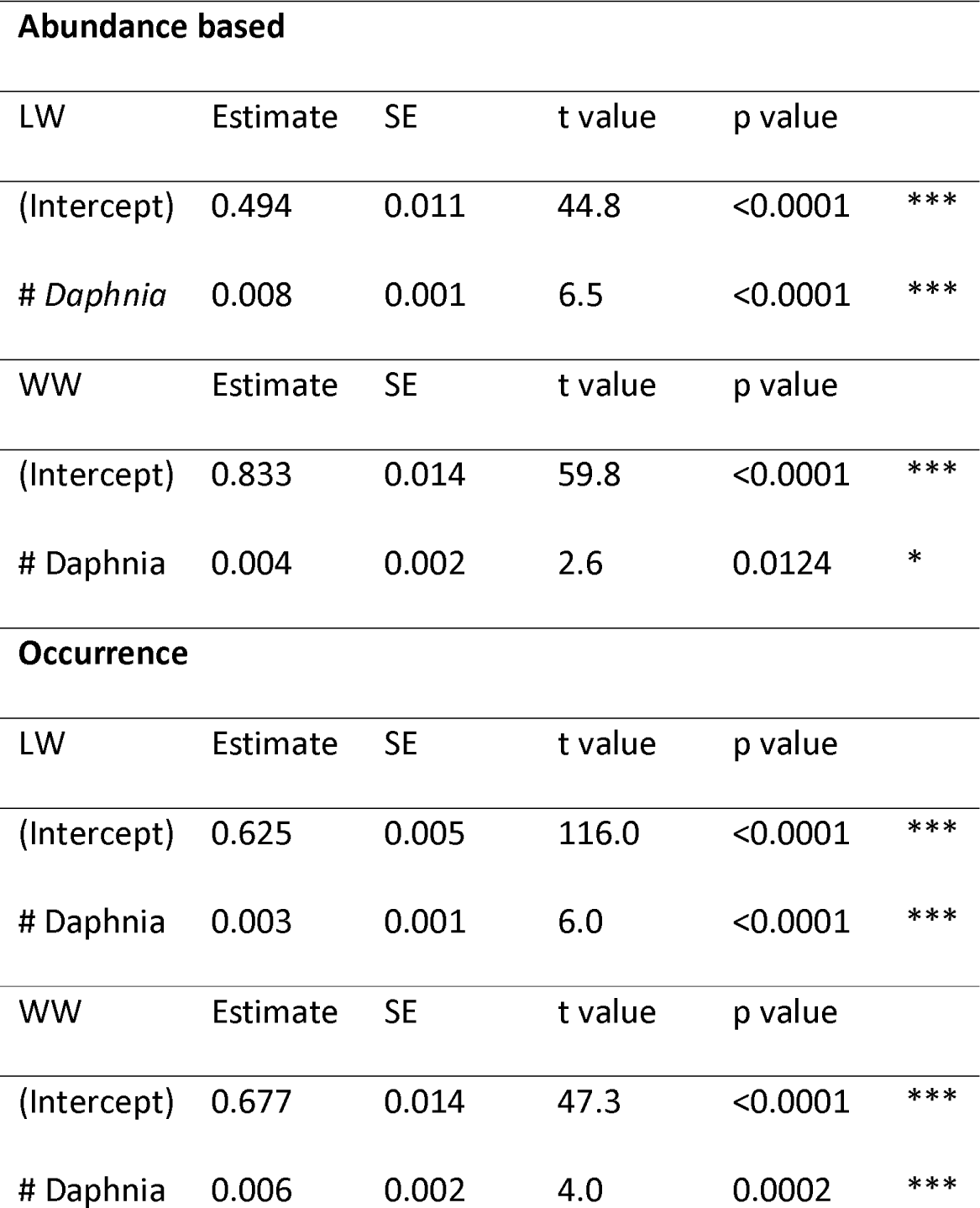
Results from the linear models testing the effect of the number of *Daphnia* on the abundance of the distance of the microbial community to the original community of lake water (LW) and waste water treatment plant effluent water (WW) 100% lake water (LW) or 10% WWTP effluent water (WW) and 90% LW for both abundance-based and occurrence based Bray-Curtis dissimilarity.

The complete output of the AncomBC analysis is presented in supplementary file 1. Of a total of 25 genera that were negatively correlated with *Daphnia* abundance, nine were characteristic of lake water, seven were neutral and nine originated from WWTP (Figure 6).

**Figure 6:**
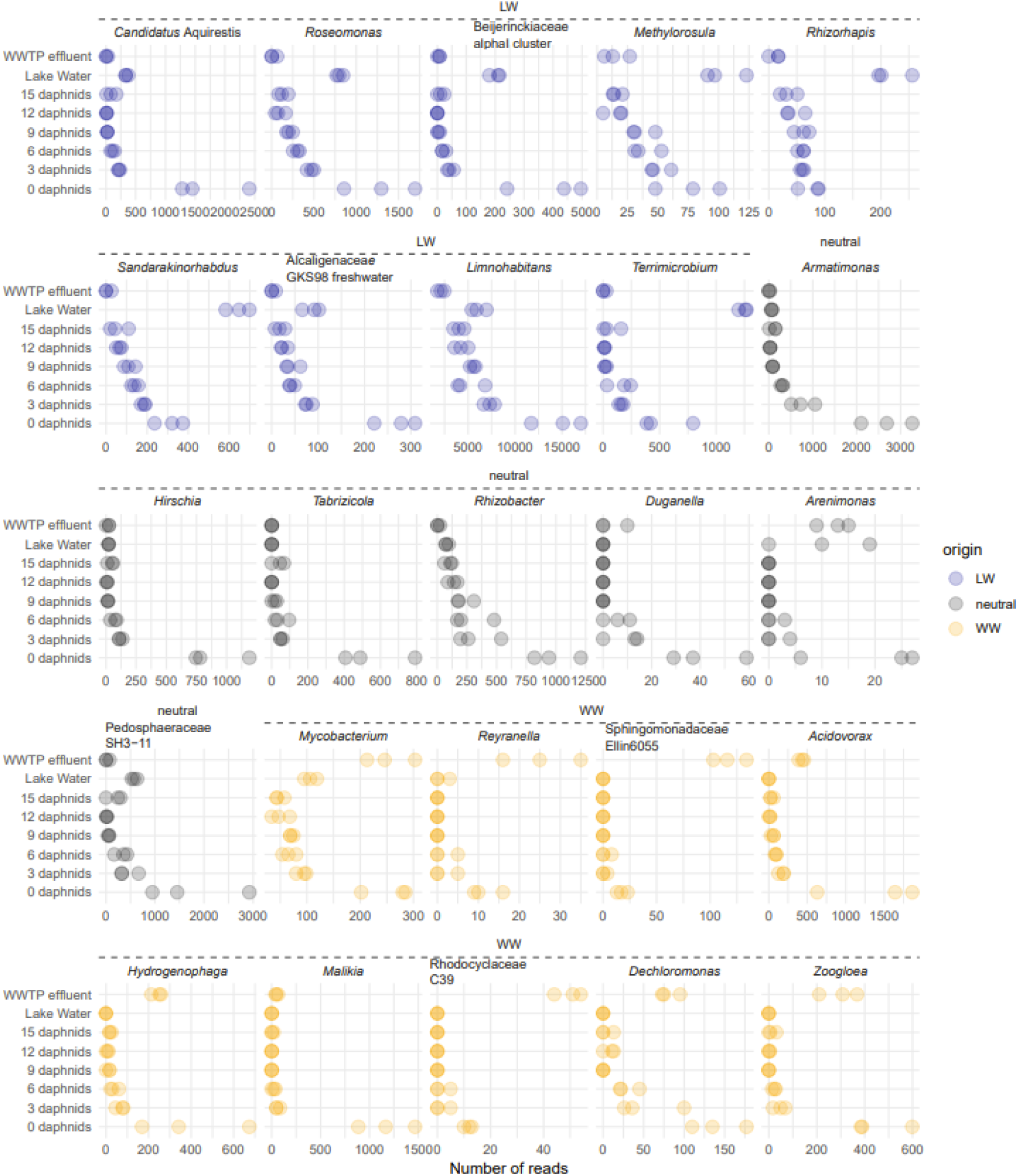
Number of reads for genera that are significantly negatively correlated with the number of daphnids in the treatments. The colours of the points denote whether a specific genus was significantly overrepresented in waste water effluent water (black), lake water (blue) or neither (orange) according to AncomBC analysis.

## Discussion

The analysis of different size fractions of the Lake Maggiore microbial community strongly suggests a particle-bound life-style for *intI*1-harbouring bacteria. This is evidenced through a two orders of magnitude higher abundance of this gene in the >1.2 µm size fraction, representing particle-associated bacteria, than in the >0.2 µm size fraction, representing free-living bacteria. Such a result provides support to the formerly suggested attachment of *intI*1-harbouring bacteria to sinking particles, such as small organic particles or lake snow (Grossart and Simon 1993). Generally, *intI*1 was temporarily enriched, and no difference was seen between the two sampling stations, which might suggest specific growth of certain *intI*1-containing bacteria within the system, as compared to continuous seeding in from WWTP effluents.

In the fraction above 50 µm, that we defined as zooplankton-associated bacteria, in addition to bacteria potentially associated to large biofilms on very large particles, we could only observe sporadic spikes of the *intI*1 gene. This strongly suggests a sudden intense enrichment of the zooplankton microbiota with specific *intI*1-hosting bacteria. Zooplankton microbiota composition is considered to be very flexible (Eckert et al. 2021) and such short-term enrichments of zooplankton microbiota were observed for *e.g.*, *E. coli*, and even natural antibiotic resistance gene containing bacteria (*tet*A) and *intI*1 (Eckert et al. 2016; Di Cesare et al. 2018, 2022a). However, this does not seem to be the case in our system since the addition of *Daphnia* generally decreased *intI*1 containing bacteria even in the food-web experiment where DNA extractions were done from the whole community and thus included also the bacteria associated with the animal itself.

In the water *Daphnia* clearly had a linear impact on the removal of *intI*1-harbouring bacteria. Interestingly, no grazing effect on the relative abundance of *intI*1-harboring bacteria was observed for other grazers, such as the heterotrophic nanoflagellate *Poteriochromonas* sp. or *Rotaria macrua* , both of which feed on bacteria (Ricci 1984; Corno 2006). However, the size fraction of bacterial removal might be very different: Nano flagellates target medium sized unicellular bacteria as preferential prey (Corno and Jürgens 2008). The size range of particles that are preferentially ingested by *R. macrura* is yet to be studied, but other bdelloid rotifers with similar feeding mechanisms and head-size (*i.e.*, *Philodina* sp.) were also shown to preferentially ingest smaller sized particles in the size range of single cell bacteria (Lapinski and Tunnacliffe 2003). On the contrary *Daphnia* feeding predominantly removes larger bacteria such as filaments and/or aggregates (Langenheder and Jürgens 2001). Evidence of this is also given in the here presented 16S rRNA sequencing data where the obligate filamentous Candidatus genus *Aquirestis* was strongly removed in the presence of *Daphnia* (Hahn and Schauer 2007). Moreover, our experiment does not suggest that *Daphnia* filter feeding reduced *intI*1 containing bacteria through a cascading food web effects (Zöllner et al. 2003) but through direct removal of the bacteria themselves, which again hints to a predominant particle attached or aggregated life-style.

Since the addition of 10% WWTP effluent increased the effect of *Daphnia* on the abundance of the *intI*1 gene, we analysed the microbial community of this experiment to determine whether *Daphnia* grazing would preferentially remove WWTP derived bacteria. For this purpose, we then studied the distance the resulting communities had to the original WWTP effluent and lake water community. Distance to the original community, for both lake water and WWTP effluent, increased with increasing number of *Daphnia* abundances. In terms of occurrence-based beta diversity metrics, *Daphnia* seemed to have a stronger effect on the lake water bacteria, whereas in terms of abundance-based metrics the animals seemed to impact WWTP derived bacteria in a stronger manner. Notwithstanding these differences, it seems, however, that the effect was similar on both microbial communities. A similar pattern was seen when bacterial genera that were significantly negatively correlated to the abundance of *Daphnia* were classified as either WWTP or LW derived. Also, here important genera were categorized as deriving from either of the sources or even as neutral. The stronger effect of removal of *intI*1 might not be related to a bacterial genus in particular but simply to a preferential growth on particles of WWTP-derived bacteria compared to typical free-living pelagic bacteria from the lake.

Many of the genera that were found to be reduced by *Daphnia* are known to grow in aggregates or on surfaces, such as *Sandarakinorhabdus* aggregated with Cyanobacteria (Cai et al. 2018), *Armatimonas* growing on aquatic plants (Tamaki et al. 2011), *Hirschia,* growing on many types of surfaces (Chepkwony et al. 2019). *Reyranella* can develop biofilm and has been suggested as a potential utilizer of plastic as a carbon source (Gorokhova et al. 2021). *Acidovorax* is commonly found in wastewater (Palanisamy et al. 2022), and it is also known to be present in aquatic environments (Chen et al. 2020). *Malikia* is an important member of biofilms developed in WWTPs (Ziegler et al. 2016; Naz et al. 2018). *Dechloromonas* is particularly abundant in biological phosphorus removal systems of WWTPs (Petriglieri et al. 2021), where is prone to develop biofilm on metal surfaces (Li et al. 2020) and polymers (Shen et al. 2016). *Zoogloe* encompasses denitrifying bacteria (Zhang et al. 2023) and has been found as in important component of the algal-bacteria granular sludge (Liu et al. 2023). It can exist in aggregate form (Chakraborty et al. 2011) contributing to the development of biofilm on various polymers (Shen et al. 2013; Walczak et al. 2015; Fang et al. 2020). *Mycobacterium* is one of the bacterial genera hosting the highest number of potential pathogenic species (Bartlett et al. 2022), prone to form aggregates (Borrego et al. 2000) and develop biofilm, eventually causing infectious diseases (Esteban and García-Coca 2018). All in all, unsurprisingly, many of the removed bacteria can either be found in aggregates or even filamentous forms and are therefore preferentially ingested by *Daphnia*.

In conclusion this study shows that most of the *intI*1 hosting bacteria deriving both from wastewater and lake water grow on particulate substrates in open waters, making them particularly vulnerable to grazing by large filter feeders such as *Daphnia*. Therefore, *intI*1 containing bacteria do not comply with the definition of invasive species, but in terms of dynamics it seems that some *intI*1 containing bacteria manage to attach to particle and even grow on them, with which they seem to sink down to the sediment where the gene accumulates (Di Cesare, 2020). To which extent the gene found in deep sediments is related to viable bacteria or just DNA residues remains to be explored. This has extensive implications when considering the persistence of *intI*1 and its gene cassettes that often are ARGs in the freshwater ecosystem. The studies of *intI*1, the most promising genetic marker for pollution with determinants of antibiotic resistance in the environment, are still in their infancy and we need to increase the number of studies on long-term monitoring of various water fractions and on species interactions in order to understand the ecological context of the persistence of such a gene in surface waters.

## Acknowledgments

The author would like to thank Evelina Crippa, Gabriele Tartari, Paola Giacomotti and Arianna Orrù for technical assistance, and Acqua Novara-VCO for providing the waters of the effluent of the WWTP of Verbania. Funding was provided by *CIPAIS* (International Commission for the Protection of Italian-Swiss Waters), Program 2022-24, Section: Limnological Researches.

## Data Availability Statement

The raw amplicon reads of the bacterial community are available in the NCBI database and can be accessed under project ID PRJNA962832. All other data and R scripts can be found here https://github.com/EsterME/Daphnia_intI1.

## Author Contribution Statement

*EME, GC, ADC, GB and DF designed the study. GB, EME, GC and DF conducted the experiments. CC, RP, GB and EME collected and processed environmental samples and metadata. GB, EME and ADC conducted molecular biology, analysed the data and made graphs. The article was written by GB, EME and ADC with input from all the authors. This study was funded by grants awarded to GC, CC, RP and DF*.

## Supplementary Material

**Figure S1:**
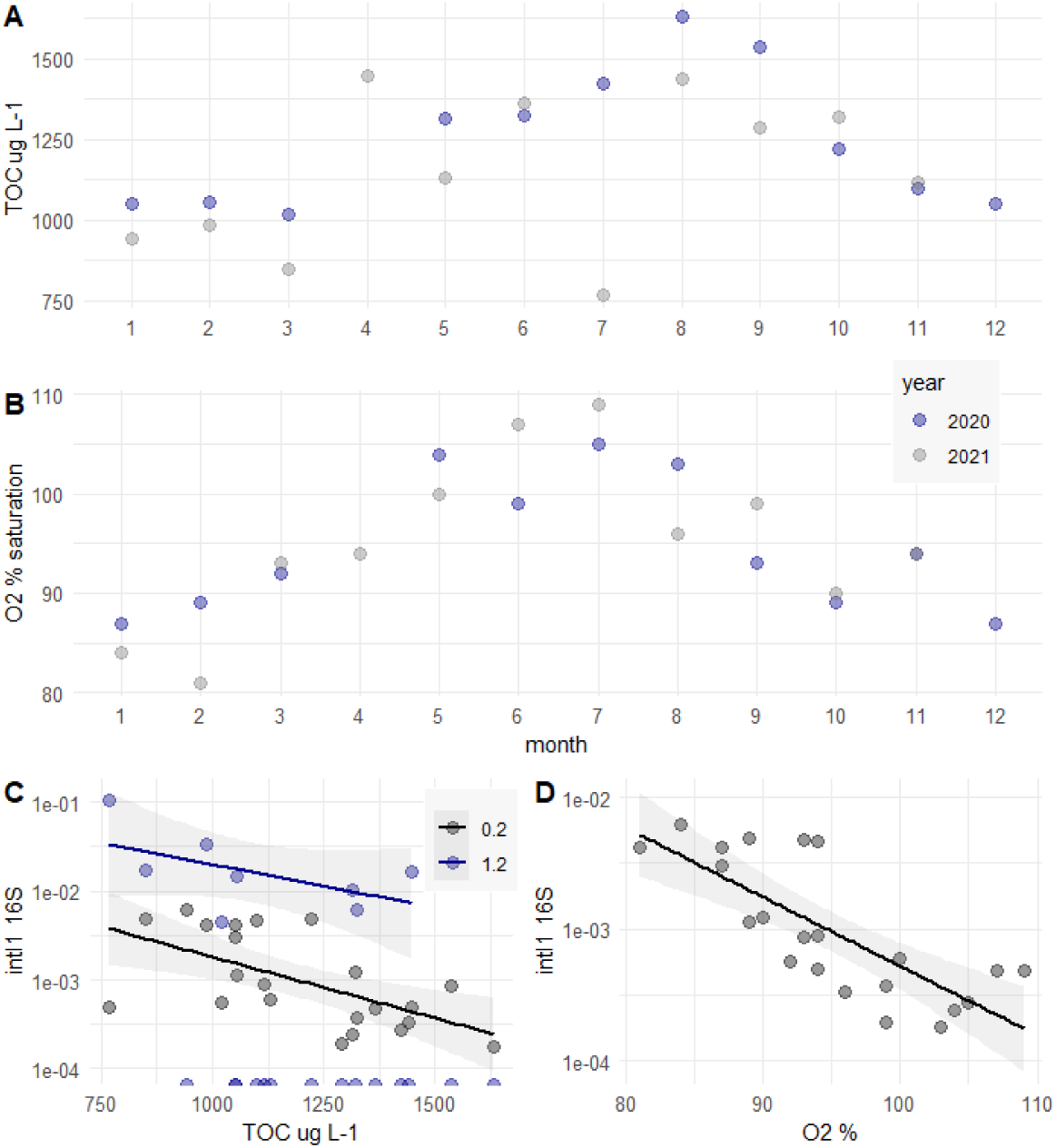
Concentration of TOC (A) and saturation of O2 (B) throughout the years 2020 and 2021 in the Euphotic zone of Lake Maggiore. Relation of relative *intI*1 abundances in the 0.2 µm and 1.2 µm fraction in the years 2020 and 2021 with total organic carbon (TOC) (C) and the saturation of % Oxygen (O2) (D).

**Table S1:**
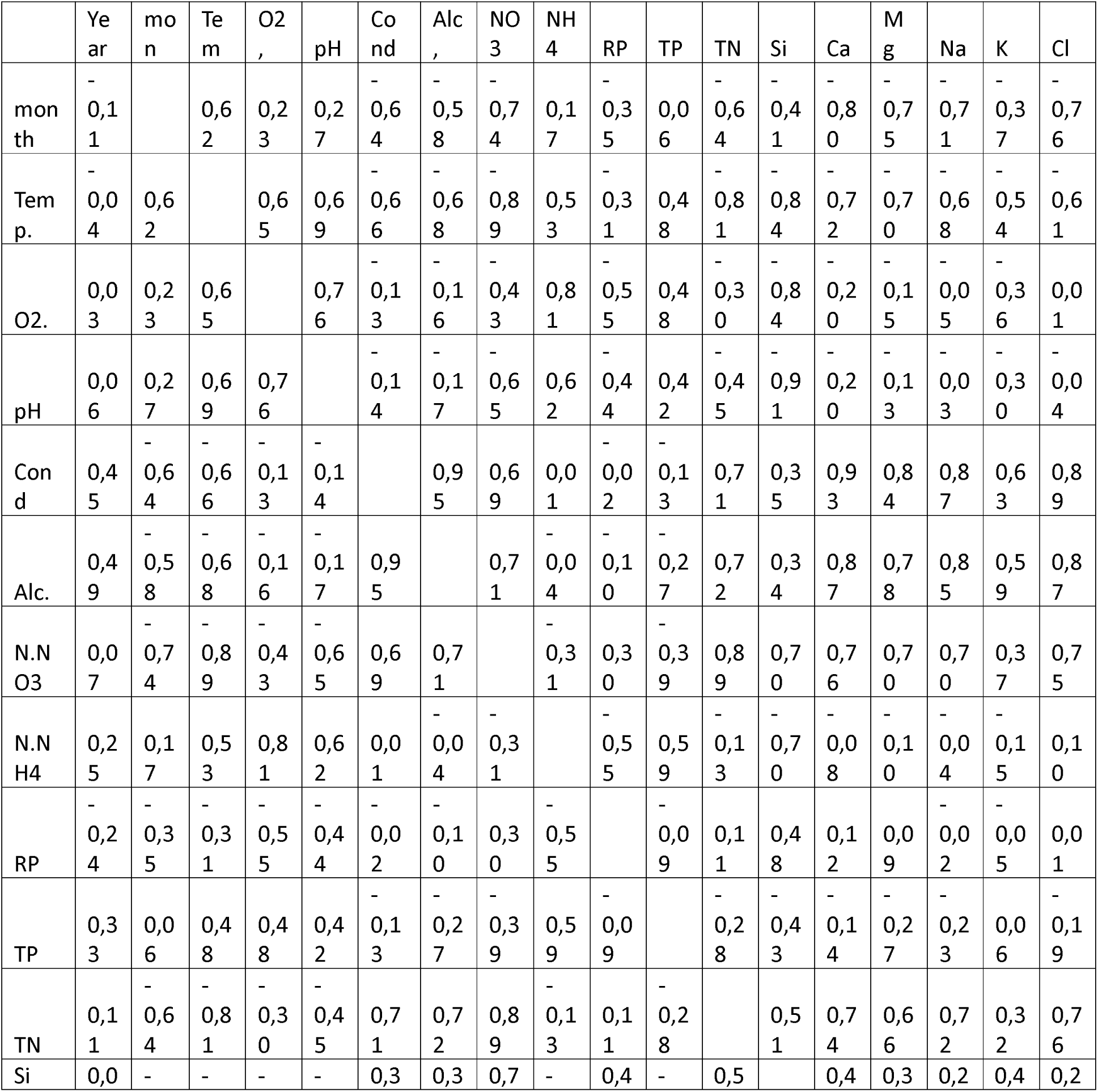
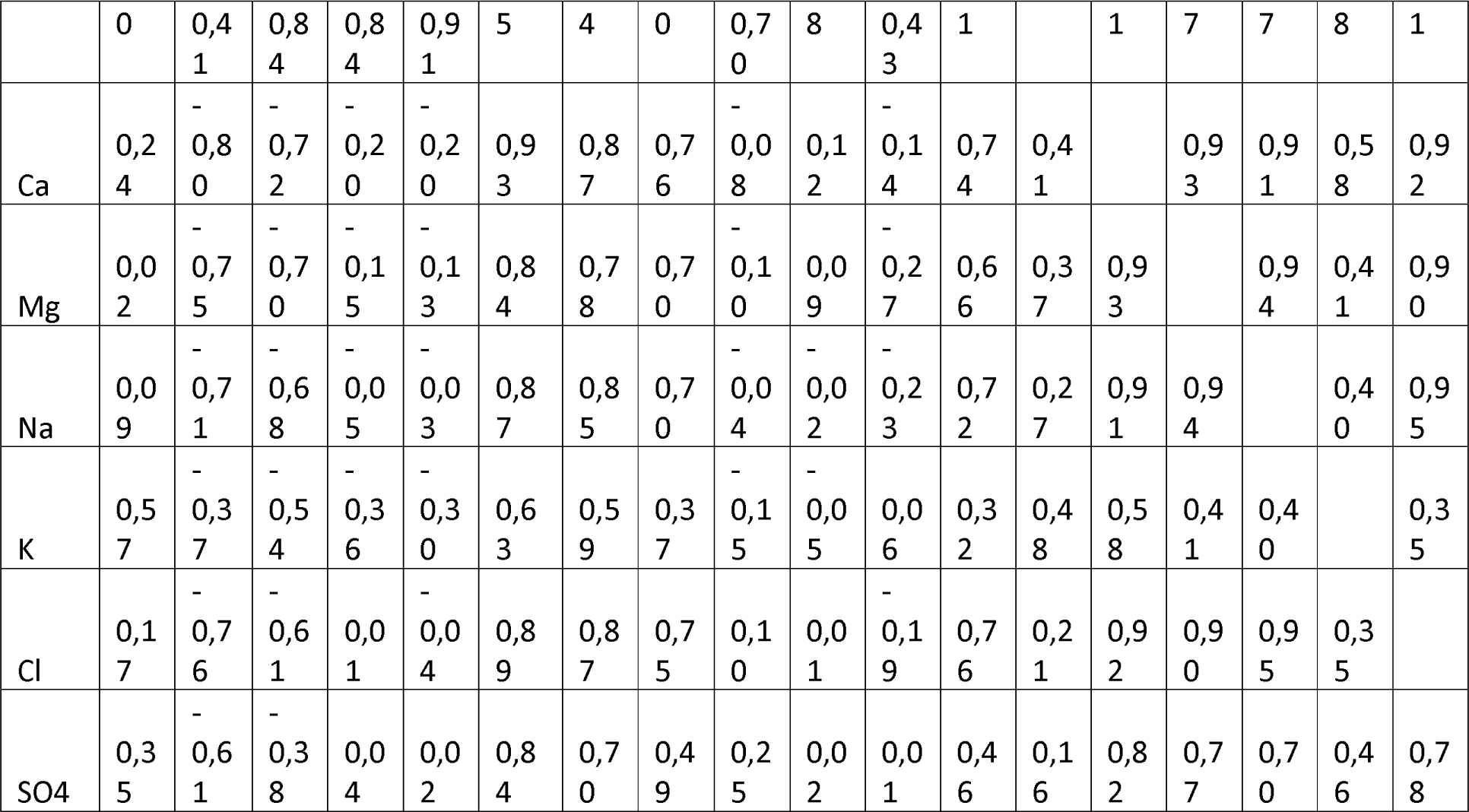
Pearson’s moment correlation between the different physic chemical parameters tested for samples from 2020 and 2021.

**Table S2:** limits of quantification of intI1 and 16S rRNA genes quantified by qPCR.

| | copies/μL per 16S rRNA gene | copies/μL per *int1* gene |
| --- | --- | --- |
| Natural bacterial community | 1548 | 11 |
| Particle associated fraction and zooplankton organisms | 50 | 225 |
| Food web experiment | 155 | 112 |
| Gradient experiment | 58 | 112 |

## References

1. Abramova, A., T. U. Berendonk, and J. Bengtsson-Palme. 2022. Meta-analysis reveals the global picture of antibiotic resistance gene prevalence across environments. 2022.01.29.478248. doi:10.1101/2022.01.29.478248

2. Barraud, O., M. C. Baclet, F. Denis, and M. C. Ploy. 2010. Quantitative multiplex real-time PCR for detecting class 1, 2 and 3 integrons. Journal of Antimicrobial Chemotherapy 65: 1642–1645. doi:10.1093/jac/dkq167

3. Bartlett, A., D. Padfield, L. Lear, R. Bendall, and M. Vos. 2022. A comprehensive list of bacterial pathogens infecting humans. Microbiology 168: 001269. doi:10.1099/mic.0.001269

4. Bertoni, R., C. Callieri, G. Corno, S. Rasconi, E. Caravati, and M. Contesini. 2010. Long-term trends of epilimnetic and hypolimnetic bacteria and organic carbon in a deep holo-oligomictic lake. Hydrobiologia 644: 279–287. doi:10.1007/s10750-010-0150-x

5. Borrego, S., E. Niubó, O. Ancheta, and M. E. Espinosa. 2000. Study of the microbial aggregation inmycobacterium using image analysis and electron microscopy. Tissue and Cell 32: 494–500. doi:10.1016/S0040-8166(00)80005-1

6. Bustin, S. A., V. Benes, J. A. Garson, and others. 2009. The MIQE Guidelines: Minimum Information for Publication of Quantitative Real-Time PCR Experiments. Clinical Chemistry 55: 611–622. doi:10.1373/clinchem.2008.112797

7. Cacace, D., D. Fatta-Kassinos, C. M. Manaia, and others. 2019. Antibiotic resistance genes in treated wastewater and in the receiving water bodies: A pan-European survey of urban settings. Water Research 162: 320–330. doi:10.1016/j.watres.2019.06.039

8. Cai, H., H. Cui, Y. Zeng, M. An, and H. Jiang. 2018. Sandarakinorhabdus cyanobacteriorum sp. nov., a novel bacterium isolated from cyanobacterial aggregates in a eutrophic lake. International Journal of Systematic and Evolutionary Microbiology 68: 730–735.

9. Callahan, B. J., P. J. McMurdie, M. J. Rosen, A. W. Han, A. J. A. Johnson, and S. P. Holmes. 2016. DADA2: High-resolution sample inference from Illumina amplicon data. Nat Methods 13: 581–583. doi:10.1038/nmeth.3869

10. Callieri, C., R. Sabatino, A. D. Cesare, and R. Bertoni. 2022. Spatial-temporal study of cluster 5 picocyanobacteria and exopolymeric microgels in Lake Maggiore. Advances in Oceanography and Limnology 13. doi:10.4081/aiol.2022.11043

11. Chakraborty, A., E. E. Roden, J. Schieber, and F. Picardal. 2011. Enhanced Growth of Acidovorax sp. Strain 2AN during Nitrate-Dependent Fe(II) Oxidation in Batch and Continuous-Flow Systems. Applied and Environmental Microbiology 77: 8548–8556. doi:10.1128/AEM.06214-11

12. Chaturvedi, P., A. Singh, P. Chowdhary, A. Pandey, and P. Gupta. 2021. Occurrence of emerging sulfonamide resistance (sul1 and sul2) associated with mobile integrons-integrase (intI1 and intI2) in riverine systems. Science of The Total Environment 751: 142217. doi:10.1016/j.scitotenv.2020.142217

13. Chen, L., J. Zhang, H. Dai, B. X. Hu, J. Tong, D. Gui, X. Zhang, and C. Xia. 2020. Comparison of the groundwater microbial community in a salt-freshwater mixing zone during the dry and wet seasons. Journal of Environmental Management 271: 110969. doi:10.1016/j.jenvman.2020.110969

14. Chepkwony, N. K., C. Berne, and Y. V. Brun. 2019. Comparative analysis of ionic strength tolerance between freshwater and marine Caulobacterales adhesins. Journal of Bacteriology 201: 10–1128.

15. Claus O. Wilke. 2020. cowplot: Streamlined Plot Theme and Plot Annotations for “ggplot2.”

16. Corno, G. 2006. Effects of nutrient availability and Ochromonas sp. predation on size and composition of a simplified aquatic bacterial community. FEMS Microbiology Ecology 58: 354–363. doi:10.1111/j.1574-6941.2006.00185.x

17. Corno, G., T. Ghaly, R. Sabatino, E. M. Eckert, S. Galafassi, M. R. Gillings, and A. Di Cesare. 2023. Class 1 integron and related antimicrobial resistance gene dynamics along a complex freshwater system affected by different anthropogenic pressures. Environmental Pollution 316: 120601. doi:10.1016/j.envpol.2022.120601

18. Corno, G., and K. Jürgens. 2006. Direct and Indirect Effects of Protist Predation on Population Size Structure of a Bacterial Strain with High Phenotypic Plasticity. Applied and Environmental Microbiology 72: 78–86. doi:10.1128/AEM.72.1.78-86.2006

19. Corno, G., and K. Jürgens. 2008. Structural and functional patterns of bacterial communities in response to protist predation along an experimental productivity gradient. Environmental Microbiology 10: 2857–2871. doi:10.1111/j.1462-2920.2008.01713.x

20. De Bernardi, R., G. Giussani, M. Manca, T. Ruffoni, and A. Savia. 1985. Laboratory effects of three species of Daphnia on Scenedesmus population growth and on selected environmental parameters. SIL Proceedings, 1922-2010 22: 3030–3034. doi:10.1080/03680770.1983.11897827

21. Di Cesare, A., E. M. Eckert, A. Teruggi, D. Fontaneto, R. Bertoni, C. Callieri, and G. Corno. 2015. Constitutive presence of antibiotic resistance genes within the bacterial community of a large subalpine lake. Mol Ecol 24: 3888–3900. doi:10.1111/mec.13293

22. Di Cesare, A., D. Fontaneto, J. Doppelbauer, and G. Corno. 2016. Fitness and Recovery of Bacterial Communities and Antibiotic Resistance Genes in Urban Wastewaters Exposed to Classical Disinfection Treatments. Environ. Sci. Technol. 50: 10153–10161. doi:10.1021/acs.est.6b02268

23. Di Cesare, A., G. M. Luna, C. Vignaroli, S. Pasquaroli, S. Tota, P. Paroncini, and F. Biavasco. 2013. Aquaculture Can Promote the Presence and Spread of Antibiotic-Resistant Enterococci in Marine Sediments. PLOS ONE 8: e62838. doi:10.1371/journal.pone.0062838

24. Di Cesare, A., S. Petrin, D. Fontaneto, and others. 2018. ddPCR applied on archived Continuous Plankton Recorder samples reveals long-term occurrence of class 1 integrons and a sulphonamide resistance gene in marine plankton communities. Environmental microbiology reports 10: 458–464.

25. Di Cesare, A., F. Riva, N. Colinas, and others. 2022a. Zooplankton as a Transitional Host for Escherichia coli in Freshwater. Applied and Environmental Microbiology 88: e02522–21.

26. Di Cesare, A., R. Sabatino, Y. Yang, D. Brambilla, P. Li, D. Fontaneto, E. M. Eckert, and G. Corno. 2022b. Contribution of plasmidome, metal resistome and integrases to the persistence of the antibiotic resistome in aquatic environments. Environmental Pollution 297: 118774. doi:10.1016/j.envpol.2021.118774

27. Di Cesare, A., E. M. Eckert, C. Cottin, A. Bouchez, C. Callieri, M. Cortesini, A. Lami, and G. Corno. 2020. The vertical distribution of tetA and intI1 in a deep lake is rather due to sedimentation than to resuspension. FEMS Microbiology Ecology 96: fiaa002. doi:10.1093/femsec/fiaa002

28. Ebert. 1993. The trade-off between offspring size and number in Daphnia magna: the influence of genetic, environmental and maternal effects.

29. Eckert, E. M., N. Anicic, and D. Fontaneto. 2021. Freshwater zooplankton microbiome composition is highly flexible and strongly influenced by the environment. Molecular Ecology 30: 1545–1558. doi:10.1111/mec.15815

30. Eckert, E. M., A. Di Cesare, B. Stenzel, D. Fontaneto, and G. Corno. 2016. Daphnia as a refuge for an antibiotic resistance gene in an experimental freshwater community. Science of the Total Environment 571: 77–81.

31. Esteban, J., and M. García-Coca. 2018. Mycobacterium Biofilms. Frontiers in Microbiology 8.

32. Fang, D., A. Wu, L. Huang, Q. Shen, Q. Zhang, L. Jiang, and F. Ji. 2020. Polymer substrate reshapes the microbial assemblage and metabolic patterns within a biofilm denitrification system. Chemical Engineering Journal 387: 124128. doi:10.1016/j.cej.2020.124128

33. Gillings, M. R. 2014. Integrons: past, present, and future. Microbiol Mol Biol Rev 78: 257–277. doi:10.1128/MMBR.00056-13

34. Gillings, M. R. 2017. Class 1 integrons as invasive species. Current Opinion in Microbiology 38: 10–15. doi:10.1016/j.mib.2017.03.002

35. Gillings, M. R., W. H. Gaze, A. Pruden, K. Smalla, J. M. Tiedje, and Y.-G. Zhu. 2015. Using the class 1 integron-integrase gene as a proxy for anthropogenic pollution. ISME J 9: 1269–1279. doi:10.1038/ismej.2014.226

36. Gorokhova, E., A. Motiei, and R. El-Shehawy. 2021. Understanding Biofilm Formation in Ecotoxicological Assays With Natural and Anthropogenic Particulates. Frontiers in Microbiology 12.

37. Grossart, H.-P., and M. Simon. 1993. Limnetic macroscopic organic aggregates (lake snow): Occurrence, characteristics, and microbial dynamics in Lake Constance. Limnology and Oceanography 38: 532– 546. doi:10.4319/lo.1993.38.3.0532

38. Haenelt, S., G. Wang, J. C. Kasmanas, F. Musat, H. H. Richnow, U. N. da Rocha, J. A. Müller, and N. Musat. 2023. The fate of sulfonamide resistance genes and anthropogenic pollution marker intI1 after discharge of wastewater into a pristine river stream. Frontiers in Microbiology 14.

39. Hahn, M. W., and M. Schauer. 2007. ‘Candidatus Aquirestis calciphila’and ‘Candidatus Haliscomenobacter calcifugiens’, filamentous, planktonic bacteria inhabiting natural lakes. International journal of systematic and evolutionary microbiology 57: 936–940.

40. Herlemann, D. P., M. Labrenz, K. Jürgens, S. Bertilsson, J. J. Waniek, and A. F. Andersson. 2011. Transitions in bacterial communities along the 2000 km salinity gradient of the Baltic Sea. ISME J 5: 1571–1579. doi:10.1038/ismej.2011.41

41. Korajkic, A., P. Wanjugi, L. Brooks, Y. Cao, and V. J. Harwood. 2019. Persistence and Decay of Fecal Microbiota in Aquatic Habitats. Microbiology and Molecular Biology Reviews 83: 10.1128/mmbr.00005-19. doi:10.1128/mmbr.00005-19

42. LaMontagne, J. m., and E. McCauley. 2001. Maternal effects in Daphnia: what mothers are telling their offspring and do they listen? Ecology Letters 4: 64–71. doi:10.1046/j.1461-0248.2001.00197.x

43. Lampert, W. 2001. Survival in a varying environment phenotypic and genotypic responses in Daphnia populations. Limnetica 20: 3–14. doi:10.23818/limn.20.02

44. Langenheder, S., and K. Jürgens. 2001. Regulation of bacterial biomass and community structure by metazoan and protozoan predation. Limnology and Oceanography 46: 121–134. doi:10.4319/lo.2001.46.1.0121

45. Lapinski, J., and A. Tunnacliffe. 2003. Reduction of suspended biomass in municipal wastewater using bdelloid rotifers. Water Research 37: 2027–2034. doi:10.1016/S0043-1354(02)00626-7

46. Lee, J., F. Ju, A. Maile-Moskowitz, and others. 2021. Unraveling the riverine antibiotic resistome: The downstream fate of anthropogenic inputs. Water Research 197: 117050. doi:10.1016/j.watres.2021.117050

47. Lenth R. 2023. Lenth R (2023). _emmeans: Estimated Marginal Means, aka Least-Squares Means_. R package version 1.8.4-1.

48. Li, W., Q. Tan, W. Zhou, J. Chen, Y. Li, F. Wang, and J. Zhang. 2020. Impact of substrate material and chlorine/chloramine on the composition and function of a young biofilm microbial community as revealed by high-throughput 16S rRNA sequencing. Chemosphere 242 125310. doi:10.1016/j.chemosphere.2019.125310

49. Lin, H., M. Eggesbø, and S. D. Peddada 2022. Linear and nonlinear correlation estimators unveil undescribed taxa interactions in microbiome data. Nat Commun 13: 4946. doi:10.1038/s41467-022-32243-x

50. Liu, Z., J. Wang, S. Zhang, and others. 2023. Formation characteristics of algal-bacteria granular sludge under low-light environment: From sludge characteristics, extracellular polymeric substances to microbial community. Bioresource Technology 376: 128851. doi:10.1016/j.biortech.2023.128851

51. Lüdecke, D., M. Ben-Shachar, I. Patil, P. Waggoner, and D. Makowski. 2021. performance: An R Package for Assessment, Comparison and Testing of Statistical Models. JOSS 6: 3139. doi:10.21105/joss.03139

52. Marano, R. B. M., T. Fernandes, C. M. Manaia, and others. 2020. A global multinational survey of cefotaxime-resistant coliforms in urban wastewater treatment plants. Environment International 144: 106035. doi:10.1016/j.envint.2020.106035

53. McMurdie, P. J., and S. Holmes. 2013. phyloseq: An R Package for Reproducible Interactive Analysis and Graphics of Microbiome Census Data. PLOS ONE 8: e61217. doi:10.1371/journal.pone.0061217

54. Munk, P., C. Brinch, F. D. Møller, and others. 2022. Genomic analysis of sewage from 101 countries reveals global landscape of antimicrobial resistance. Nat Commun 13: 7251. doi:10.1038/s41467-022-34312-7

55. Naz, I., D. Hodgson, A. Smith, and others. 2018. Investigation of the active biofilm communities on polypropylene filter media in a fixed biofilm reactor for wastewater treatment. Journal of Chemical Technology & Biotechnology 93: 3264–3275. doi:10.1002/jctb.5686

56. Pagès, H., P. Aboyoun, R. Gentleman, and others. 2023. Biostrings: Efficient manipulation of biological strings.doi:10.18129/B9.bioc.Biostrings

57. Palanisamy, V., V. Gajendiran, and K. Mani. 2022. Meta-analysis to identify the core microbiome in diverse wastewater. Int. J. Environ. Sci. Technol. 19: 5079–5096. doi:10.1007/s13762-021-03349-4

58. Pereyra, P. J. 2016. Revisiting the use of the invasive species concept: An empirical approach. Austral Ecology 41: 519–528. doi:10.1111/aec.12340

59. Petriglieri, F., C. Singleton, M. Peces, J. F. Petersen, M. Nierychlo, and P. H. Nielsen. 2021. “Candidatus Dechloromonas phosphoritropha” and “Ca. D. phosphorivorans”, novel polyphosphate accumulating organisms abundant in wastewater treatment systems. ISME J 15: 3605–3614. doi:10.1038/s41396-021-01029-2

60. Quast, C., E. Pruesse, P. Yilmaz, J. Gerken, T. Schweer, P. Yarza, J. Peplies, and F. O. Glöckner. 2013. The SILVA ribosomal RNA gene database project: improved data processing and web-based tools. Nucleic Acids Research 41: D590–D596. doi:10.1093/nar/gks1219

61. R Core Team. 2022. R: A language and environment for statistical computing. R Foundation for Statistical Computing, Vienna, Austria.

62. Ricci, C. 1984. Culturing of some bdelloid rotifers. Hydrobiologia 112: 45–51. doi:10.1007/BF00007665

63. Rogora, M., M. Austoni, R. Caroni, and others. 2021a. Temporal changes in nutrients in a deep oligomictic lake: the role of external loads versus climate change. J Limnol 80. doi:10.4081/jlimnol.2021.2051

64. Rogora, M., M. Austoni, R. Caroni, and others. 2021b. Temporal changes in nutrients in a deep oligomictic lake: the role of external loads versus internal processes. Journal of Limnology 80. doi:10.4081/jlimnol.2021.2051

65. Shen, Z., Y. Yin, and J. Wang. 2016. Biological denitrification using poly(butanediol succinate) as electron donor. Appl Microbiol Biotechnol 100: 6047–6053. doi:10.1007/s00253-016-7435-6

66. Shen, Z., Y. Zhou, J. Hu, and J. Wang. 2013. Denitrification performance and microbial diversity in a packed-bed bioreactor using biodegradable polymer as carbon source and biofilm support. Journal of Hazardous Materials 250–251: 431–438. doi:10.1016/j.jhazmat.2013.02.026

67. Subirats, J., A. Di Cesare, S. Varela della Giustina, A. Fiorentino, E. M. Eckert, S. Rodriguez-Mozaz, C. M. Borrego, and G. Corno. 2019. High-quality treated wastewater causes remarkable changes in natural microbial communities and intI1 gene abundance. Water Research 167: 114895. doi:10.1016/j.watres.2019.114895

68. Tamaki, H., Y. Tanaka, H. Matsuzawa, M. Muramatsu, X.-Y. Meng, S. Hanada, K. Mori, and Y. Kamagata. 2011. Armatimonas rosea gen. nov., sp. nov., of a novel bacterial phylum, Armatimonadetes phyl. nov., formally called the candidate phylum OP10. International journal of systematic and evolutionary microbiology 61: 1442–1447.

69. Walczak, M., M. Swiontek Brzezinska, A. Sionkowska, M. Michalska, U. Jankiewicz, and E. Deja-Sikora. 2015. Biofilm formation on the surface of polylactide during its biodegradation in different environments. Colloids and Surfaces B: Biointerfaces 136: 340–345. doi:10.1016/j.colsurfb.2015.09.036

70. Wickham H. 2016. ggplot2: Elegant Graphics for Data Analysis. Springer-Verlag New York, 2016.

71. Zhang, J., Y. Shao, Z. Li, G. Han, X. Jing, N. Wang, J. Xu, and G. Chen. 2023. Characteristics analysis of plastisphere biofilm and effect of aging products on nitrogen metabolizing flora in microcosm wetlands experiment. Journal of Hazardous Materials 452: 131336. doi:10.1016/j.jhazmat.2023.131336

72. Zheng, W., J. Huyan, Z. Tian, Y. Zhang, and X. Wen. 2020. Clinical class 1 integron-integrase gene – A promising indicator to monitor the abundance and elimination of antibiotic resistance genes in an urban wastewater treatment plant. Environment International 135: 105372. doi:10.1016/j.envint.2019.105372

73. Ziegler, A. S., S. J. McIlroy, P. Larsen, M. Albertsen, A. A. Hansen, N. Heinen, and P. H. Nielsen. 2016. Dynamics of the Fouling Layer Microbial Community in a Membrane Bioreactor. PLOS ONE 11: e0158811. doi:10.1371/journal.pone.0158811

74. Zolkefli, N., S. S. Sharuddin, M. Z. M. Yusoff, M. A. Hassan, T. Maeda, and N. Ramli. 2020. A Review of Current and Emerging Approaches for Water Pollution Monitoring. Water 12: 3417. doi:10.3390/w12123417

75. Zöllner, E., B. Santer, M. Boersma, H. Hoppe, and K. Jürgens. 2003. Cascading predation effects of Daphnia and copepods on microbial food web components. Freshwater Biology 48: 2174–2193.

