## Supplementary Figures for "Zooplankton impacts on the persistence of the anthropogenic pollution marker *intI*1 in lake water"

**Supplementary Material**

**Table S1**: Pearson’s moment correlation between the different physic chemical parameters tested for samples from 2020 and 2021.

|  | Year | mon | Tem | O2, | pH | Cond | Alc, | NO3 | NH4 | RP | TP | TN | Si | Ca | Mg | Na | K | Cl |
| --- | --- | --- | --- | --- | --- | --- | --- | --- | --- | --- | --- | --- | --- | --- | --- | --- | --- | --- |
| month | -0,11 |  | 0,62 | 0,23 | 0,27 | -0,64 | -0,58 | -0,74 | 0,17 | -0,35 | 0,06 | -0,64 | -0,41 | -0,80 | -0,75 | -0,71 | -0,37 | -0,76 |
| Temp. | -0,04 | 0,62 |  | 0,65 | 0,69 | -0,66 | -0,68 | -0,89 | 0,53 | -0,31 | 0,48 | -0,81 | -0,84 | -0,72 | -0,70 | -0,68 | -0,54 | -0,61 |
| O2. | 0,03 | 0,23 | 0,65 |  | 0,76 | -0,13 | -0,16 | -0,43 | 0,81 | -0,55 | 0,48 | -0,30 | -0,84 | -0,20 | -0,15 | -0,05 | -0,36 | 0,01 |
| pH | 0,06 | 0,27 | 0,69 | 0,76 |  | -0,14 | -0,17 | -0,65 | 0,62 | -0,44 | 0,42 | -0,45 | -0,91 | -0,20 | -0,13 | -0,03 | -0,30 | -0,04 |
| Cond | 0,45 | -0,64 | -0,66 | -0,13 | -0,14 |  | 0,95 | 0,69 | 0,01 | -0,02 | -0,13 | 0,71 | 0,35 | 0,93 | 0,84 | 0,87 | 0,63 | 0,89 |
| Alc. | 0,49 | -0,58 | -0,68 | -0,16 | -0,17 | 0,95 |  | 0,71 | -0,04 | -0,10 | -0,27 | 0,72 | 0,34 | 0,87 | 0,78 | 0,85 | 0,59 | 0,87 |
| N.NO3 | 0,07 | -0,74 | -0,89 | -0,43 | -0,65 | 0,69 | 0,71 |  | -0,31 | 0,30 | -0,39 | 0,89 | 0,70 | 0,76 | 0,70 | 0,70 | 0,37 | 0,75 |
| N.NH4 | 0,25 | 0,17 | 0,53 | 0,81 | 0,62 | 0,01 | -0,04 | -0,31 |  | -0,55 | 0,59 | -0,13 | -0,70 | -0,08 | -0,10 | -0,04 | -0,15 | 0,10 |
| RP | -0,24 | -0,35 | -0,31 | -0,55 | -0,44 | -0,02 | -0,10 | 0,30 | -0,55 |  | -0,09 | 0,11 | 0,48 | 0,12 | 0,09 | -0,02 | -0,05 | 0,01 |
| TP | 0,33 | 0,06 | 0,48 | 0,48 | 0,42 | -0,13 | -0,27 | -0,39 | 0,59 | -0,09 |  | -0,28 | -0,43 | -0,14 | -0,27 | -0,23 | 0,06 | -0,19 |
| TN | 0,11 | -0,64 | -0,81 | -0,30 | -0,45 | 0,71 | 0,72 | 0,89 | -0,13 | 0,11 | -0,28 |  | 0,51 | 0,74 | 0,66 | 0,72 | 0,32 | 0,76 |
| Si | 0,00 | -0,41 | -0,84 | -0,84 | -0,91 | 0,35 | 0,34 | 0,70 | -0,70 | 0,48 | -0,43 | 0,51 |  | 0,41 | 0,37 | 0,27 | 0,48 | 0,21 |
| Ca | 0,24 | -0,80 | -0,72 | -0,20 | -0,20 | 0,93 | 0,87 | 0,76 | -0,08 | 0,12 | -0,14 | 0,74 | 0,41 |  | 0,93 | 0,91 | 0,58 | 0,92 |
| Mg | 0,02 | -0,75 | -0,70 | -0,15 | -0,13 | 0,84 | 0,78 | 0,70 | -0,10 | 0,09 | -0,27 | 0,66 | 0,37 | 0,93 |  | 0,94 | 0,41 | 0,90 |
| Na | 0,09 | -0,71 | -0,68 | -0,05 | -0,03 | 0,87 | 0,85 | 0,70 | -0,04 | -0,02 | -0,23 | 0,72 | 0,27 | 0,91 | 0,94 |  | 0,40 | 0,95 |
| K | 0,57 | -0,37 | -0,54 | -0,36 | -0,30 | 0,63 | 0,59 | 0,37 | -0,15 | -0,05 | 0,06 | 0,32 | 0,48 | 0,58 | 0,41 | 0,40 |  | 0,35 |
| Cl | 0,17 | -0,76 | -0,61 | 0,01 | -0,04 | 0,89 | 0,87 | 0,75 | 0,10 | 0,01 | -0,19 | 0,76 | 0,21 | 0,92 | 0,90 | 0,95 | 0,35 |  |
| SO4 | 0,35 | -0,61 | -0,38 | 0,04 | 0,02 | 0,84 | 0,70 | 0,49 | 0,25 | 0,02 | 0,01 | 0,46 | 0,16 | 0,82 | 0,77 | 0,70 | 0,46 | 0,78 |

**Table S2**: limits of quantification of intI1 and 16S rRNA genes quantified by qPCR.

|  | **copies/μL per 16S rRNA gene** | **copies/μL per *intI*1 gene** |
| --- | --- | --- |
| Natural bacterial community | 1548 | 11 |
| Particle associated fraction and zooplankton organisms | 50 | 225 |
| Food web experiment | 155 | 112 |
| Gradient experiment | 58 | 112 |

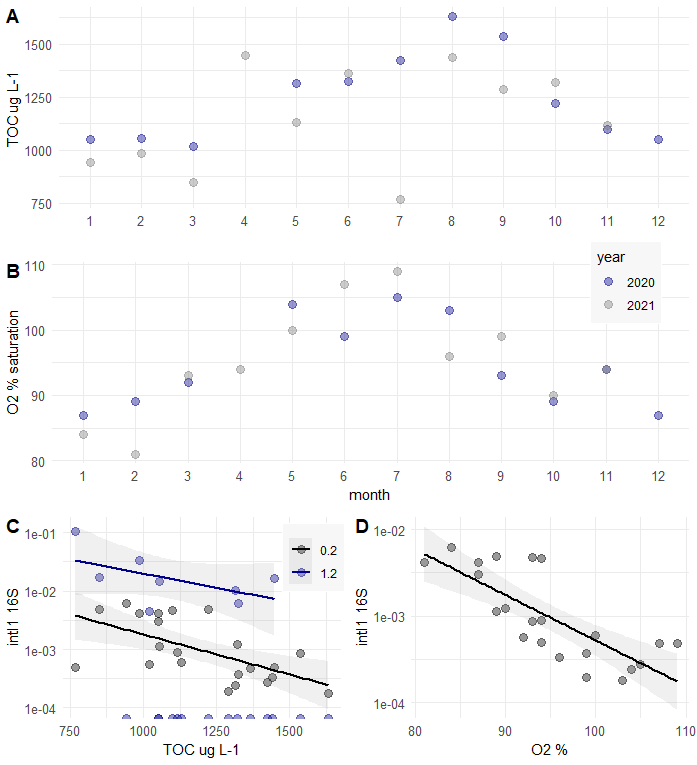

**Figure S1**:Concentration of TOC (A) and saturation of O2 (B) throughout the years 2020 and 2021 in the Euphotic zone of Lake Maggiore. Relation of relative *intI*1 abundances in the 0.2 µm and 1.2 µm fraction in the years 2020 and 2021 with total organic carbon (TOC) (C) and the saturation of % Oxygen (O2) (D).

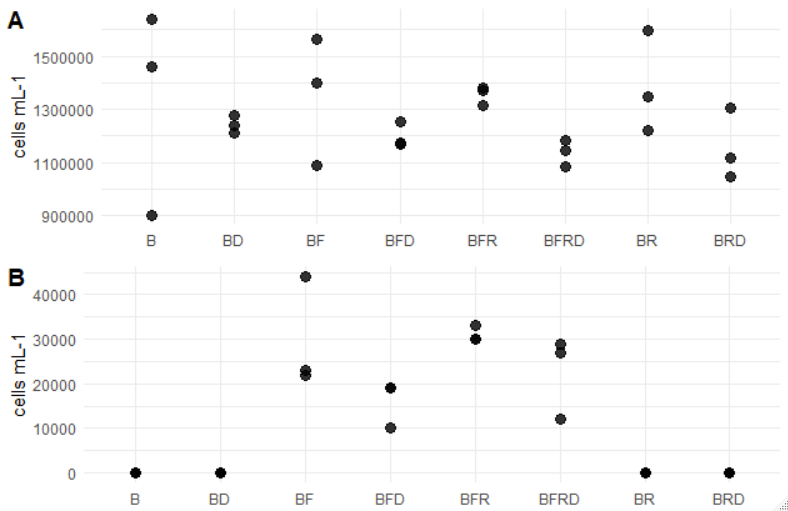

**Figure S2**: Cells per ml in the food web experiment as measured by Flow Cytometry for A) Bacteria and B) Flagellates. Food web composition is denoted as follows: B=Bacteria, D=Daphnia, F=Flagellate, R=Rotifer.
